# Accumulation and maintenance of information in evolution

**DOI:** 10.1101/2021.12.23.473971

**Authors:** Michal Hledík, Nick Barton, Gašper Tkačik

## Abstract

Selection accumulates information in the genome — it guides stochastically evolving populations towards states (genotype frequencies) that would be unlikely under neutrality. This can be quantified as the Kullback-Leibler (KL) divergence between the actual distribution of genotype frequencies and the corresponding neutral distribution. First, we show that this population-level information sets an upper bound on the information at the level of genotype and phenotype, limiting how precisely they can be specified by selection. Next, we study how the accumulation and maintenance of information is limited by the cost of selection, measured as the genetic load or the relative fitness variance, both of which we connect to the control-theoretic KL cost of control. The information accumulation rate is upper bounded by the population size times the cost of selection. This bound is very general, and applies across models (Wright-Fisher, Moran, diffusion) and to arbitrary forms of selection, mutation and recombination. Finally, the cost of maintaining information depends on how it is encoded: specifying a single allele out of two is expensive, but one bit encoded among many weakly specified loci (as in a polygenic trait) is cheap.

## 1 Introduction

Throughout evolution, selection accumulates information in the genome. It guides evolving populations towards fitter phenotypes, genotypes and genotype frequencies, which would be highly unlikely to arise by chance. This information – the degree to which selection can control the stochastic process of evolution – has been a long-standing subject of research [1]–[7], and relates to basic questions in evolutionary biology and genetics.

### How well can selection specify the genotype and the phenotype?

The degree to which within- and between-species genetic variation are shaped by selection has been the subject of the neutralist-selectionist debate [8]–[11]. Today, we know that much of the human genome is involved in various biochemical processes [12], [13], but this does not mean that it is strongly shaped by selection [14]–[16]. Here we ask a related question in information-theoretic terms: how much information can selection accumulate and maintain in the genome? Much of the sequence is to some degree random, and given its size *l* ≈ 3 × 10^9^ base pairs, it likely contains far less information than the maximum conceivable 6 × 10^9^ bits of information. A similar question has been raised in the context of origin of life: given high mutation rates, how much information could be maintained in the genome of early organisms [2]?

Analogous questions can be asked about the phenotype. How many traits can selection optimize? It is easy to list a large number of potentially relevant traits: take the expression of all genes in all cell types and conditions, or regulatory interactions between pairs of genes. For a fit organism, these traits need to be specified with some precision, and this precision is likely limited (even if it is to some degree facilitated by correlations among traits). For example, a study of selective constraint on human gene expression [17] gave evidence of constraint, but overall, this seems weak. Given the large number of possibly important phenotypes, how precisely can selection specify them?

### Quantifying genetic information

An established method in bioinformatics quantifies the information content of a short genomic motif, such as a binding site, by comparing an alignment of its instances across the genome to the genomic background [18], [19]. Our definition of genetic information is mathematically similar, but aims to apply more generally (to large regions without multiple instances available). It is therefore based in theoretical population genetics rather than sequence data analysis. A key related concept is the repeatability of evolution [20], [21]. Evolution is stochastic due to genetic drift and mutation, but selection can reduce the space of possible outcomes. For example, suppose that in a sequence of length *l, n* sites are under strong selection for specific nucleotides. By fixing those nucleotides, selection will accumulate 2*n* bits of information. Meanwhile, the remaining *l* – *n* sites will be occupied by random nucleotides, and if a replicate population evolves under identical conditions, the *l* – *n* nucleotides will likely be different. Therefore our concept of information in a sequence is inversely related to how differently it could have evolved under identical conditions.

In general, however, the information content of the genome cannot be quantified by simply counting the sites that are under selection. A single bit of information can be spread across many loci under weak selection – a phenomenon particularly relevant when selection acts on polygenic traits, long recognized in quantitative genetics and described by the infinitesimal model [22], [23]. Polygenicity and weak selection also resolve the apparent contradiction between the variety of phenotypes, or biochemical processes involving the DNA, and the lack of strong selective constraint on all of them. Selection might act on a small number of high level traits, which are influenced by large numbers of loci spread across the genome (described by the omnigenic model [24]), which experience only weak selection individually.

In Section 2, we define information on three levels – the population state (genotype frequencies), the genotype, and the phenotype. There are simple inequalities between the three levels. This means that the upper bound on information accumulation rate, which we prove at the population level, also implies a bound at the genotype and phenotype levels. We use the KL divergence, a central quantity in information theory [25] to quantify the difference between their actual distribution and their corresponding neutral distribution.

Notably, the neutral phenotype distribution corresponds approximately to the phenotype distribution among random DNA sequences. Recent work with random mutant libraries suggests that for some phenotypes, this distribution is accessible experimentally (gene expression driven by random promoters [26]–[28] or enhancers [29]). Any departure from this neutral distribution amounts to accumulation of information.

### Cost of information

After defining what genetic information means, we ask how quickly it can accumulate and how much of it can be maintained. We look for answers in terms of the cost of selection – the amount of relative fitness variation in a population. This cost, traditionally measured as the relative fitness variance or the genetic load, is itself limited. In a population with constant size, relative fitness is proportional to the expected number of offspring, and the number of offspring can only vary between zero and the reproductive capacity of the organism.

We rely on an information-theoretic measure of cost of selection, which is itself upper bounded by the relative fitness variance and genetic load, but has favorable mathematical properties. It relates the cost of selection to the KL cost of control [30]–[32], or the thermodynamic power [33].

The relationship between information accumulation rate and the cost of selection has been studied by Kimura [1] and later Worden [3], MacKay [4] and Barton [7]. In Sec. 3 we discuss these works in more detail and present a new, more general bound. The problem of maintenance has been studied by Eigen [2], Watkins [5] and Peck and Waxman [6]. We discuss these in Sec. 4 and present example calculations that suggest general trends in the amount of information that can be maintained per unit cost.

## 2 Quantifying genetic information

The measures of information studied in this paper are based on comparisons between the distributions of various variables under selection versus neutrality. The focus on probability distributions accounts for the stochasticity of evolution, and the difference between the distributions with and without selection corresponds to the control that selection exerts on evolution. We quantify this difference in bits, using the KL divergence [25]

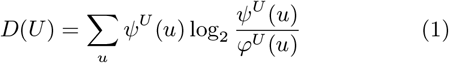

where *U* is a variable that takes values *u* with probabilities *ψ^U^* (*u*) with selection and *φ^u^* (*u*) under neutrality. Below we focus on three variables – genotype frequencies (which describe population states), genotypes and phenotypes.

For a pair of variables *U, V*, statistical dependencies are reflected in their joint and conditional KL divergence, *D*(*U,V*) and *D*(*U*|*V*) (see SI Sec. S1 for the definitions). Both are nonnegative quantities, and they follow the chain rule

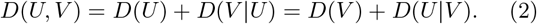

The chain rule allows a comparison of the effects of selection on different variables, as well as on the same variable at different times.

### 2.1 Population-level information

Evolution is a stochastic process happening to populations, and genotype frequencies form the state space. We use *X* to denote the genotype frequencies as a random variable, with each value *x* being a vector with an element *x_g_* for each genotype *g*, normalized as ∑_*g*_ *x_g_* = 1. As an example, Fig. 1A shows a common evolutionary scenario where a single locus, two allele system starts from a single copy of a beneficial allele *A*, and later the frequency evolves stochastically.

**Figure 1:**
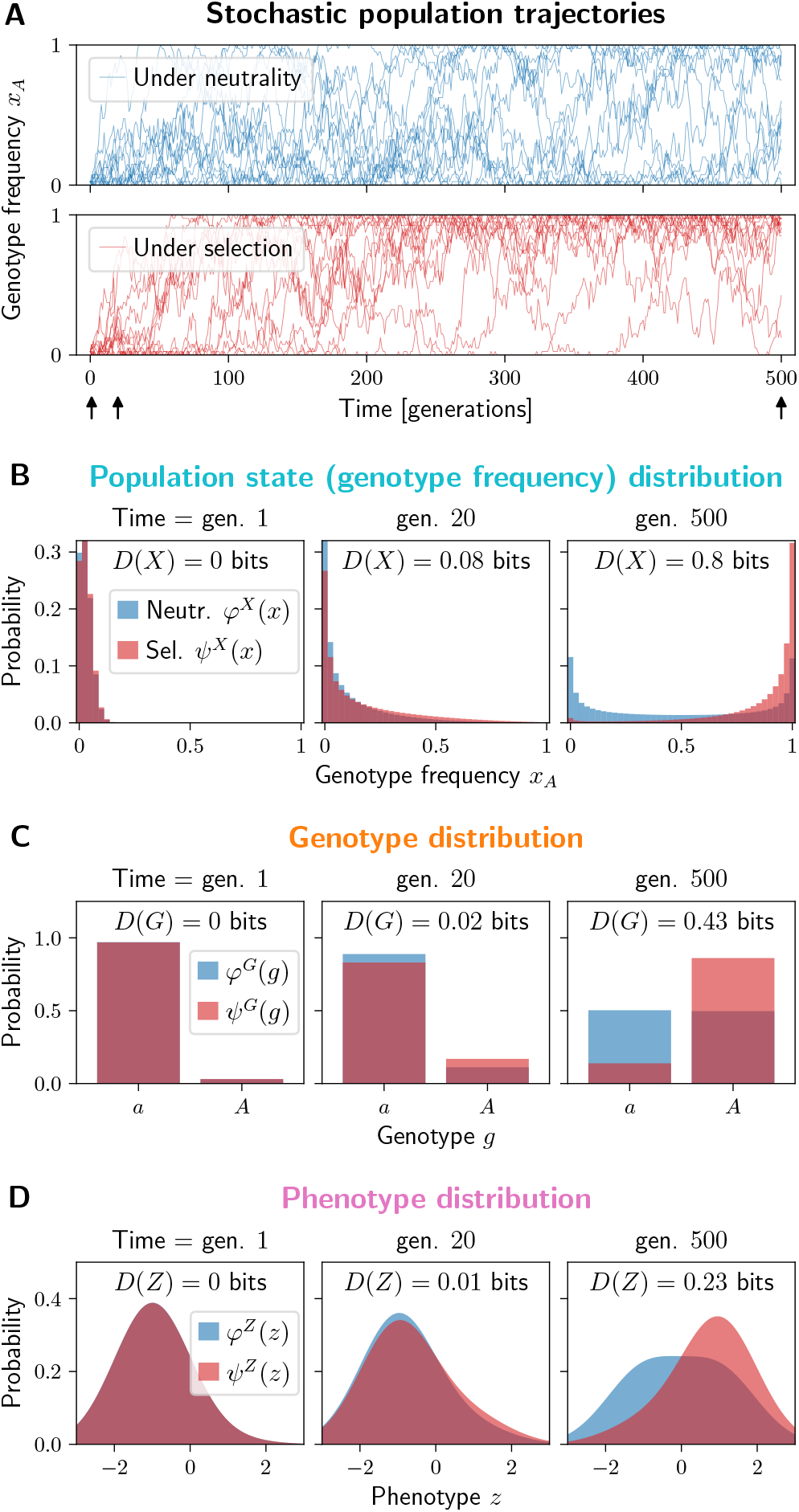
Selection controls the evolution of a single locus, two allele system and drives the distribution of the population states, genotype and phenotype away from neutrality. (A) Stochastic trajectories of the frequency *x_A_* of the beneficial allele *A*, under neutrality and under selection (blue and red). The allele A starts at a single copy, and under selection it tends to increase in frequency. Black arrows indicate the times when the distributions are plotted in (B-D). At time = 500 generations, the system is approximately stationary. (B-D) The probability distributions of the genotype frequency *x_A_* (B), genotype *g* (C) and a noisy phenotype *z* (D) under neutrality (blue) and under selection (red) after a varying number of generations of evolution. The associated measures of information *D*(*X*), *D*(*G*) and *D*(*Z*) are indicated in each panel. (B) The neutral distribution *φ^X^* converges to a symmetric U shape, while the distribution under selection is biased towards high frequencies of the beneficial allele *A*. The information *D*(*X*) increases over time. (C) The neutral genotype distribution *φ^G^* converges to a uniform distribution, due to symmetry between alleles *a* and *A*. Under selection (*ψ^G^*), the beneficial allele *A* has a higher probability, but it does not dominate completely, so the genotype-level information *D*(*G*) is less than the maximum 1 bit. *D*(*G*) is also upper bounded by *D*(*X*). (D) A phenotype with different means and a Gaussian noise for each allele, 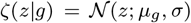 with *μ_a_* = −1, *μA* = +1 and *σ* =1. The information *D*(*Z*) is upper bounded by *D*(*G*), with a gap due to the partially overlapping distributions *ζ*(*z*|*a*) and *ζ*(*z*|*A*). Generated using a haploid Wright-Fisher model (SI Sec. S4) with population size *N* = 40, mutation rate *μ* = 0.005 and fitness 1 (allele *a*) and 1.05 (allele *A*).

*X* takes values *x* with probabilities *ψ^X^* (*x*) under selection and *φ^X^* (*x*) under neutrality. Fig. 1B shows examples of these distributions for the single locus system at three different times. In general, these distributions are shaped by various evolutionary forces – mutation, drift, recombination, selection (*ψ^X^* only), and others. We refer to *D*(*X*), the KL divergence between *ψ^X^* and *φ^X^*, as the population-level information.

The example in Fig. 1 illustrates two important phenomena we discuss in the rest of the paper. The first phenomenon is the accumulation of information. A population evolves from an initial distribution (in the simplest case, *ψ^X^* = *φ^X^* and *D*(*X*) = 0, but this is not necessary). For example, the initial state *x* may be completely specified as in Fig. 1A, or both *ψ^X^* and *φ^X^* may start at the neutral stationary distribution. Over time, selection causes *ψ^X^* to diverge from *φ^X^* and the information *D*(*X*) accumulates (Fig. 1B). We study this in detail in Sec. 3. The second phenomenon is the maintenance of information, and it takes place when both *ψ^X^* (*x*) and *φ^X^* (*x*) are stationary, and the information *D*(*X*) is constant. In Sec. 4 we study how much information can be maintained at a given cost of selection.

The population-level information *D*(*X*) has been studied under different names and in different roles [7], [34]–[36]. It captures any departure of the genotype frequency distribution *ψ^X^* from its neutral counterpart *φ^X^* – notably, selection can favor not only high frequencies of fit genotypes, but also higher or (more typically) lower amounts of genetic variation within populations. Note that *D*(*X*) refers to the effects of selection on the genotype frequencies, rather than allele frequencies. It therefore includes effects of selection on correlations between loci (linkage disequilibrium), which are generated by physical linkage, by chance in finite populations, or due to functional interactions (epistasis) – see also SI Sec. S2.

Notably, *D*(*X*) (or *D*(*G*) introduced below) appears as a term in free fitness – a quantity analogous to free energy which, under some assumptions, increases over time [35], [37], [38]. This implies that evolution maximizes the expected log-fitness while constraining *D*(*X*) – see SI Sec. S8.

### 2.2 Genotype-level information

If we sample a random genotype from a population in a given state *x*, we find the genotype *g* with a probability given simply by its frequency *ψ*^*G*|*X*^(*g*|*x*) = *φ*^*G*|*X*^(*g*|*x*) = *x_g_*. Taking into account evolutionary stochasticity, we average over all population states *x* with their probabilities *φ^X^* (*x*) or *ψ^X^* (*x*),

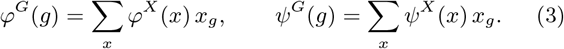

Under symmetric point mutations, the neutral distribution *φ^G^* converges to a uniform distribution over all genotypes, while selection typically concentrates *ψ^G^* among a smaller number of fit genotypes. This is also the case for the single locus system in Fig. 1C. The divergence between *ψ^G^* and *φ^G^* is the genotype-level information *D*(*G*).

If selection precisely specifies *n* out of *l* nucleotides in the genome – i.e. *ψ^G^* (*g*) is uniform over a fraction 1/4^*n*^ out of 4^*l*^ possible genotypes – this implies *D*(*G*) = 2*n* bits. This corresponds to the intuition of 2*n* bits of information encoded in the genome. More typically, selection will specify many sites only weakly (biasing the probability towards some alleles, see also Fig. 1C), and may contribute to *D*(*G*) through LD – correlations between linked or epistatically interacting sites. Without linkage or epistasis, *D*(*G*) is approximately additive across loci (see Fig. S1).

*D*(*G*) generalizes some previous definitions of genetic information [1], [3], [6] which focused on strong selection or uniform distributions, and coincides with others in important special cases [4], [5].

### 2.3 Phenotype-level information

Finally, selection controls evolution on the level of the phenotype *Z*. *Z* could be a categorical trait such as the presence/absence of a disease or the correct/incorrect protein fold, a quantitative trait, a comprehensive characterization of an individual, or its fitness. Given a genotype *g*, the probability of the phenotype *z* will be given by the possibly noisy genotype-phenotype relationship *ψ*^*Z*|*G*^ (*z*|*g*) = *φ*^*Z*|*G*^ (*z*|*g*) = *ζ*(*z*|*g*). When there are no environmental effects or intrinsic noise, *ζ*(*z*|*g*) will be concentrated at a single value z for each genotype *g*. Taking into account the variation within populations, as well as the evolutionary stochasticity, the marginal probability of *z* is

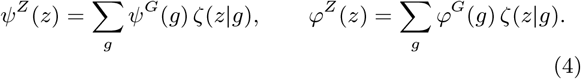

We show the distributions *ψ^Z^*, *φ^Z^* for the single locus system in Fig. 1D, where the trait has a genotype-dependent mean and a Gaussian noise. While under neutrality, *φ^Z^* tends to spread out over time, selection causes *ψ^Z^* to be more concentrated. The divergence between *ψ^Z^* and *φ^Z^* is the phenotype-level information *D*(*Z*).

If we can take the genotype distribution *φ^G^* to be uniform over all possible DNA sequences of some length, then *φ^Z^* is the phenotype distribution among such random sequences. Examples of this distribution have recently been measured experimentally, for gene expression generated by random promoter sequences in *S. cerevisiae* and *E. coli* [26], [28]. If a healthy cell requires the gene expression to be in some narrow range, this translates to a requirement on the phenotype-level information *D*(*Z*), and this requirement will increase if the expression needs to be specified across cell states.

### 2.4 The relationship between the three levels

The definitions above, combined with the chain rule (Eq. (2)) lead to a hierarchy among the three levels,

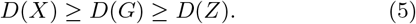

This inequality can be observed across columns of panels in Fig. 1B-D.

Intuitively, the phenotype-level information *D*(*Z*) is bounded by the genotype-level information *D*(*G*), since the information about the phenotype has to be encoded in the genome. A special case of this relationship has been noted by Worden [3], who however, worked in a deterministic setting (see SI Sec. S3). The difference between the two, *D*(*G*) – *D*(*Z*) = *D*(*G*|*Z*), can have two sources. First, the phenotype distribution *ζ*(*z*|*g*) may overlap between genotypes, causing the phenotype to be specified less precisely than the genotype (as in Fig. 1D). Second, selection may favor genotypes based on criteria other than the phenotype *Z*, such as other phenotypes or robustness.

Similarly, *D*(*G*) can only be as large as the population-level information *D*(*X*). To increase the probability of a genotype *g*, selection must increase the probability of population states with a high frequency of *g*. However, selection can also shape the patterns of genetic diversity in populations, without impacting the average genotype frequencies, therefore contributing to the difference *D*(*X*) – *D*(*G*) = *D*(*X*|*G*). In populations with weak mutation, which tend to have little diversity, this difference is small – see Fig. 2.

**Figure 2:**
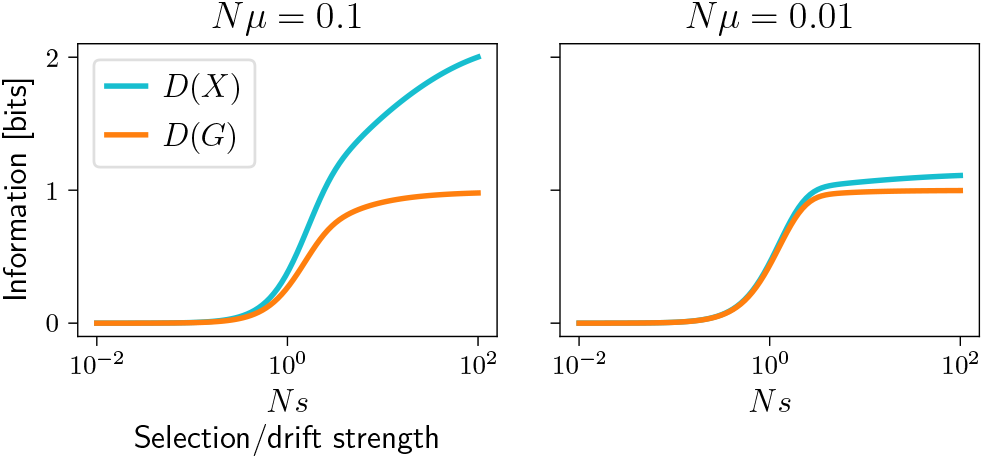
Illustration of *D*(*X*) (cyan) and *D*(*G*) (orange) for a single locus, two allele system at stationary distributions *ψ^X^*, *φ^X^* as a function of selection strength *Ns* for two different mutation strengths *Nμ*. The genotype-level information *D*(*G*) grows with *Ns*, from 0 up to 1 bit one out of the two alleles dominates, with the steepest increase around *Ns* = 1. The population-level information *D*(*X*) can be much greater than *D*(*G*) when mutation is strong, and generates diversity within the population that selection can shape (or suppress). When mutation is weak, *D*(*X*) and *D*(*G*) are similar, since the population state can be specified by the allele that is currently fixed and *D*(*X*|*G*) = 0. Computed using a Wright-Fisher model as in Fig. 1, with population size *N* = 100.

We rely on the inequalities in Eq. (5) in two ways. First, an upper bound on the population-level information *D*(*X*) which we prove in Sec. 3 also implies an upper bound on the genotype and phenotype-level information *D*(*G*) and *D*(*Z*). In other words, selection can only fine-tune the phenotype to a degree to which it can control the population state.

Second, *D*(*X*) and *D*(*G*) can be difficult to estimate directly for systems with multiple loci, due to the high dimensionality (see Fig. S1). In such situations, *D*(*Z*) for fitness or a low-dimensional phenotype *Z* can serve as a lower bound on *D*(*G*) and *D*(*X*). If *Z* is the trait under selection, or fitness itself, this lower bound can be tight. This approach is applicable even for essentially black box genotype-phenotype models, such as models of gene regulation or protein folding.

## 3 Accumulation of information

In this section we show how the rate at which *D*(*X*), the population-level information, increases over time, is limited by the population size and the variation in fitness. We start by pointing out a connection between population genetics and control theory.

### 3.1 Accumulation of information and the cost of control

We consider a population evolving over time, with a trajectory *X*^0^, *X*^1^,…, *X^T^* forming a Markov chain between generations 0 and *T* (such as in Fig. 1A). The divergence of the trajectories’ distribution from neutrality, *D*(*X*^0^, *X*^1^,…, *X^T^*), has been proposed as a measure of predictability of evolution [21]. Using the chain rule (Eq. (2)), we can decompose it in two ways,

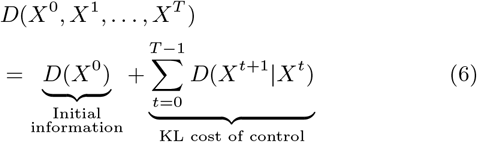

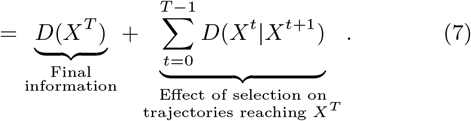

In Eq. (6), we distinguish between the divergence of the initial states *X*^0^ and the additional conditional divergence in each generation, *D*(*X*^*t*+1^|*X^t^*). The latter can be recognized as the KL cost of control, averaged over the initial states *x^t^* [30], [31]. In the context of population genetics, selection takes the role of control.

Eq. (7) makes the distinction between the distribution of endpoints *X^T^*, and the conditional distribution of the states that precede those endpoints. Selection can shape the full trajectories, but only the effects on *X^T^* constitute the final population-level information.

Together, Eq. (6) and (7) imply a bound on the information accumulated between times 0 and *T* in terms of the KL cost of control,

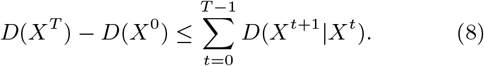

Specifically, the information accumulated over a single generation, Δ*D*(*X^t^*) = *D*(*X*^*t*+1^) – *D*(*X^t^*), is upper bounded as

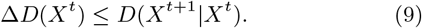

Analogous bounds for continuous time Markov chains and the diffusion approximation are in SI Sec. S6 and S7.

Note that control theory is concerned with computing optimal control policies, which maximize an imposed objective while minimizing the cost 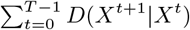. This is analogous to computing the optimal artificial selection – in fact, the KL divergence control theory framework has recently been used to study artificial selection on quantitative traits [32].

In contrast, natural selection is typically given by the biological or ecological circumstances, and not necessarily optimized in this sense. Still, the KL cost of control provides bounds on the rate at which selection accumulates information (Eq. (8,9)) and it has a meaning in population genetics, which we discuss in the next section.

We also note that Eq. (9) is related to the proof that free fitness increases over time [37], [38], see SI Sec. S8.

### 3.2 Variation in fitness as cost of control

To compute *D*(*X*^*t*+1^|*X^t^*) in population genetics, we need to specify a model. We analyze multiple general model classes in the SI: Wright-Fisher and discrete Moran models in SI Sec. S5, continuous time Moran model in SI Sec. S6 and the diffusion approximation in SI Sec. S7. In summary, the bound in Eq. (9) always takes the form

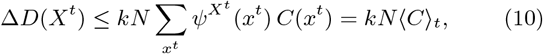

where *N* is the population size, *kN* is the number of individuals that are sampled with selection in each generation (*k* = 1 under asexual reproduction and *k* = 2 under sexual reproduction when 2 parents are sampled with selection for each individual). *C*(*x^t^*) is the cost of selection at the population state *x^t^* (see below), and 〈*C*〉_*t*_ is the expected cost at time *t*. To upper bound information accumulated over multiple generations, we need to sum over them,

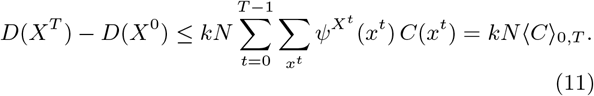

The cost *C*(*x*) is a measure of fitness variation in a population in the state *x*,

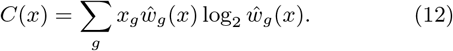

where *ŵ_g_* (*x*) is the (frequency dependent) relative fitness of genotype *g*. When sampling genotypes as parents for the next generation, *g* is picked with probability *x_g_* under neutrality and *x_g_ŵ_g_* (*x*) under selection – *C*(*x*) is the KL divergence between these two distributions.

*C*(*x*) is related to two more established measures of cost in population genetics – the relative fitness variance *V* (*x*) and the genetic load *L*(*x*), which have been studied under a number of circumstances – e.g. mutation-selection balance [39], genetic drift [40], [41], certain types of epistasis and the evolution of sex [42], [43], ongoing substitutions [44]–[46] or stabilizing selection on quantitative traits [47]. They are defined as

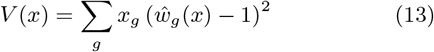

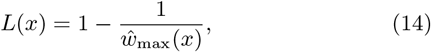

where *ŵ*_max_(*x*) is the maximum relative fitness present in the population *x, ŵ_max_*(*x*) = max_*g*;*x_g_*>0_ *ŵ_g_*(*x*). We derive the relationships between *C*(*x*), *V*(*x*) and *L*(*x*) in the SI Sec. S9. *V*(*x*) and *L*(*x*) satisfy the inequality 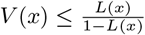 (see also [48]) and both provide an upper bound on *C*(*x*),

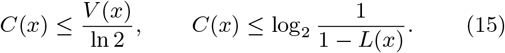

In addition, under weak selection and in the diffusion approximation, *C*(*x*) = *V*(*x*)/(2 log 2). The bounds in Eq. (10,11) can therefore also be rewritten in terms of *V*(*x*) or *L*(*x*) using Eq. (15).

Assuming constant population size, relative fitness is proportional to the expected number of offspring, and therefore limited by the species’ reproductive capacity. The quantities *ŵ*_max_(*x*), *L*(*x*), *V*(*x*) and *C*(*x*) and as a consequence Δ*D*(*X*), are therefore all limited in realistic settings (SI Sec. S9).

In the context of artificial selection or genetic algorithms, an alternative measure of cost is the population size *N*, which is the number of cultivated plants or animals, or fitness function evaluations [49], [50]. We note that according to the bounds in Eq. (10,11), the maximal accumulation rate is also proportional to *N*. Furthermore, increasing the strength of selection (and therefore *C*(*x*)) beyond an optimal value may increase the immediate response to selection, but reduces the long term response, due to loss of genetic diversity [49], [50]. Therefore in practice, *C*(*x*) will be limited even in this context.

### 3.3 Example 1: the fates of a beneficial allele

The bounds in Eq. (10,11) hold in genetically diverse populations with clonal interference or recombination. Still, it is interesting to consider the case of sequential fixation/loss of mutations, as was done previously [1], [7], [44].

Suppose that a beneficial allele *A* appears in one copy at time *t* = 0, and is guaranteed to be fixed or lost before another mutation appears that could interfere with it. The population and genotype-level information, *D*(*X^t^*) and *D*(*G^t^*), start at 0 and accumulate over time as selection tends to increase the frequency of *A* (Fig. 3A). The cumulative cost of selection *N*〈*C*〉_0,*t*_ serves as the upper bound on both *D*(*X^t^*) and *D*(*G^t^*).

**Figure 3:**
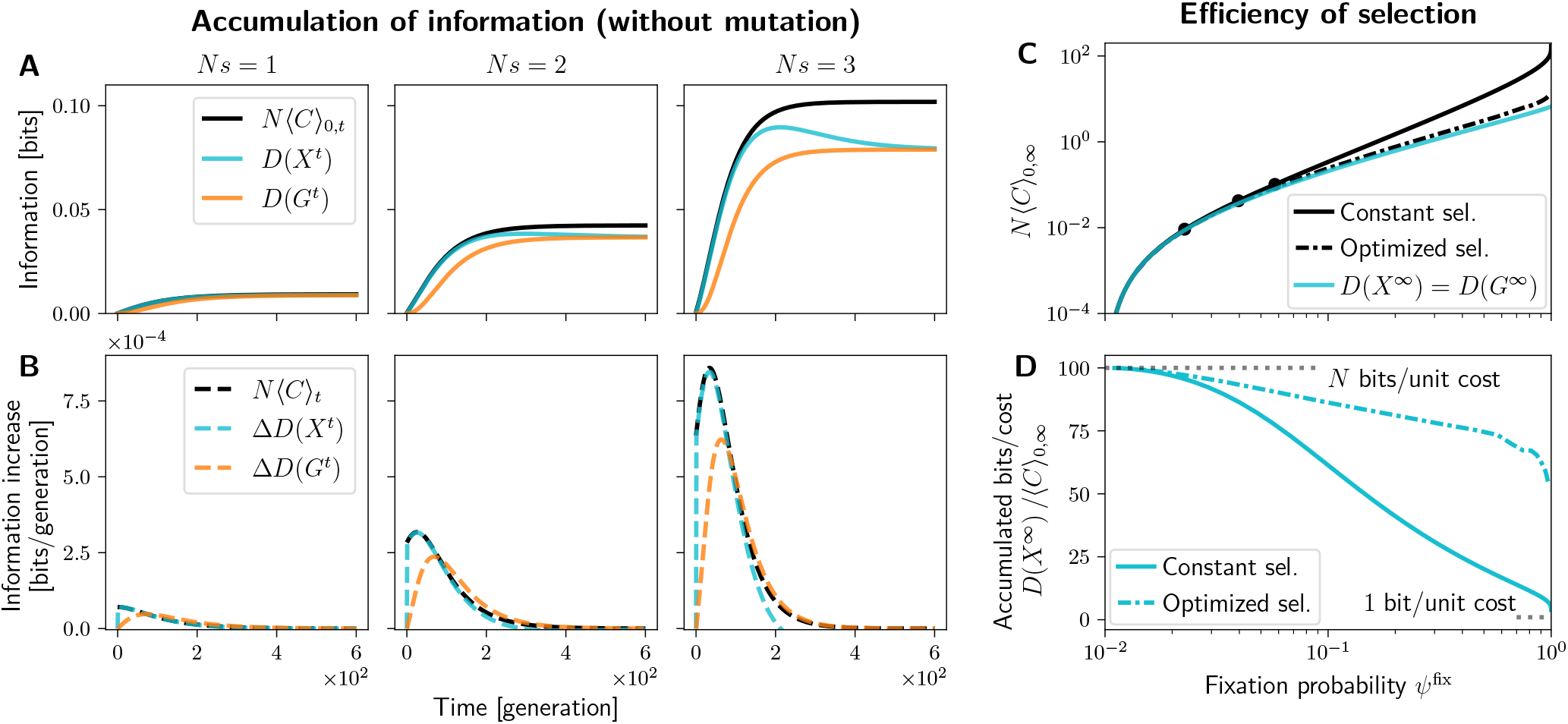
Information accumulation associated with the fixation or loss of a beneficial allele in a haploid single locus, two allele system. The beneficial allele starts at a single copy and evolves under drift and selection, but no mutation. (A) The population-level information (*D*(*X^t^*), cyan) and genotype-level information (*D*(*G^t^*), orange) over time, for three different strengths of selection (*Ns*). Both *D*(*X*^t^) and *D*(*G^t^*) start at zero, accumulate over time as selection tends to increase the frequency of the beneficial allele, and saturate as the allele is fixed or lost. The black line is the upper bound according to Eq. (11) with *k* = 1. (B) The increments in *D*(*X^t^*) and *D*(*G^t^*) per generation (cyan and orange dashed), and the upper bound according to Eq. (10) with *k* = 1 (black dashed). (C) The cyan line shows the total information accumulated, *D*(*X*^∞^) = *D*(*G*^∞^), as function of the fixation probability *ψ*^fix^. *D*(*_X_*^∞^) serves as a lower bound on *N* times the total cost of selection, plotted in black, regardless of the form selection takes. The full black line corresponds to constant selection coefficient, with black points showing the three cases in (A,B). The dash-dotted black line shows frequency dependent selection that maximizes *ψ*^fix^ (and therefore also *D*(*X*^∞^)) while constraining *N*〈*C*〉_0,∞_ (D) Same data as in (C), but the vertical axis now shows the ratio of the information *D*(*X*^∞^) and the total cost of selection 〈*C*〉_0,∞_ for constant selection (full black) and optimized frequency dependent selection (dash-dotted black). At most N bits can be accumulated per unit cost, and this is achieved at weak selection. At strong selection, this reduces to as low as 1 bit per unit cost. Figure computed using the Wright-Fisher model as in Fig. 1, with population size *N* = 100.

Note that under relatively strong selection (*Ns* = 3, Fig. 3A right), A increases in frequency considerably faster than under neutrality, leading to high *D*(*X^t^*). But some of these gains are later lost as A is fixed or lost. This is an example of how only the probabilities of endpoints, and not the shape of the trajectories, matters for the information that is ultimately accumulated (the two terms in Eq. (7)).

The increments in *D*(*X^t^*) and *D*(*G^t^*) in each generation are plotted in Fig. 3B, along with the bound by *N*〈*C*〉_*t*_, Eq. (10). The bound on Δ*D*(*X^t^*) is relatively tight. Δ*D*(*G^t^*) can temporarily exceed *N*〈*C*〉_*t*_, since the accumulation bound in Eq. (10) does not directly apply to the genotype level, but this is only a transient phenomenon due to the inequality between the cumulative genotype- and population-level information *D*(*G^t^*) ≤ *D*(*X^t^*).

Both *D*(*X^t^*) and *D*(*G^t^*) saturate at the same value *D*(*X*^∞^) = *D*(^*G*∞^), since the ultimate fate of the population is given simply by whether the allele *A* is fixed or lost. The fixation probability is 1/*N* under neutrality and *ψ*^fix^ = *ψ*^*X*∞^ (1) = *Ψ*^*G*∞^ (*A*) under selection, and the accumulated information is a function of this probability,

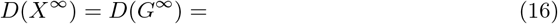

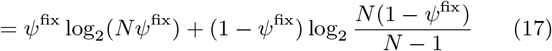

This function is plotted in cyan in Fig. 3C. According to Eq. (11), it provides a lower bound on the total cost, *N*〈*C*〉)_0,∞_ ≥ *D*(*X*^∞^), given a fixation probability. This holds when the allele A has a constant, frequency independent selective advantage, as in the three examples in Fig. 3A,B (full black line and black points in Fig. 3C). By computing a suitable frequency dependent selection, which optimizes the fixation probability while constraining the total cost *N*〈*C*〉_0_,∞, we can reduce the cost considerably (dash-dotted black line in Fig. 3C, see SI Sec. S11 and Fig. S4 for details). This is achieved by making selection weaker at high frequencies, where the risk of losing A is low. Still, the cost stays above *D*(*X*^∞^), as it has to under arbitrary frequency and time-dependent selection.

Under both forms of selection, the bound is only tight when selection is weak. To emphasize this, we plot the information accumulated per unit cost, *D*(*X*^∞^)/〈*C*〉_0,∞_, as function of the fixation probability *ψ*^fix^ in Fig. 3D. At weak selection, *ψ*^fix^ is only perturbed a little from its neutral value 1/*N*, but up to *N* bits can be accumulated per unit cost. A special case of this was shown by Barton [7]. Similar scaling with *N* was also found in a different setting by Kimura [45].

Stronger selection accumulates more information, but at a disproportionately higher cost, since a large part of it is spent on shaping trajectories rather than outcomes. In the extreme case, to achieve *ψ*^fix^ = 1, only individuals carrying the *A* allele can be allowed to reproduce and *A* gets fixed in only one generation – a highly unlikely way to fixation under neutrality. In this case, selection has the same effect on each genotype sampled as a parent in the first generation, as on the allele that is ultimately fixed (both are A with probability 1/*N* under neutrality and 1 under selection). As a result, the cost is equal to the accumulated information, 〈*C*〈_0,∞_ = *D*〈*G*^∞^) = *D*(*X*^∞^) and only 1 bit per unit cost is accumulated (Fig. 3D). This is why previous results derived in deterministic settings [1], [3] claimed much more stringent limits on accumulation of information.

### 3.4 Example 2: accumulation of information under mutation

Unlike the example above, real systems experience ongoing mutation. On the one hand, mutation is necessary to supply beneficial alleles for adaptation, but on the other hand, mutation can disrupt existing adaptation. In this section, we assume that the single locus, two allele system starts at the neutral stationary distribution with *D*(*X*^0^) = *D*(*G*^0^) = 0, and then selection is turned on. Adaptation exploits copies of the allele A that either segregate in the population by chance at time 0, or arise later by mutation.

Fig. 4A shows the information *D*(*X*^*t*^) and *D*(*G*^1^) over time. Accumulation take place on the time scale of 1/*μ*. Note that the bound Eq. (11) is not very tight. This is even more apparent in Fig. 4B, where the average cost per generation *N*〈*C*〉_*t*_ remains positive even after the system has reached the new stationary state, while the increments in *D*(*X^t^*) and *D*(*G^t^*) are zero. This corresponds to the cost of maintaining information, which we discuss in Sec. 4.

**Figure 4:**
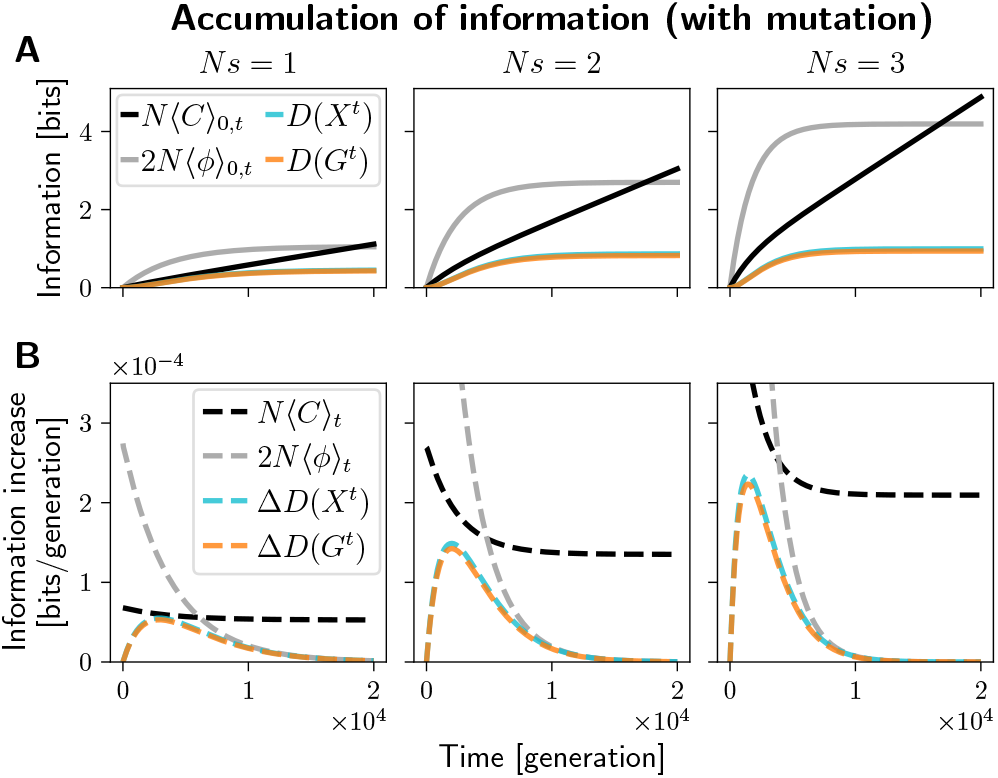
Information accumulation in a single locus, two allele system and the associated upper bounds. The system starts from a neutral stationary distribution over allele frequencies, where *D*(*X*^0^) = *D*(*G*^0^) = 0. Then it evolves under selection with varying strengths (*Ns*, left to right) for 2 × 10^4^ generations. (A) The cumulative information at the population level (*D*(*X*^0^), cyan) and genotype level (*D*(*G*^0^), orange) over time. Due to the weak mutation *Nμ* = 0.01, the two measures of information are similar. The black and gray lines show upper bounds by the cumulative cost of selection and the cumulative fitness flux. (B) The increments in information per generation, Δ*D*(*X*^t^) (cyan dashed) and Δ*D*(*G^t^*) (orange dashed) and the upper bounds on these increments in terms of the cost of selection *kN*〈*C*〉_*t*_ (black, in this case *k* = 1) and the expected fitness flux 2*N*〈*ψ*〉_*t*_ (gray). Note that the cost of selection bound is briefly nearly tight under weak selection (*Ns* = 1, left), and the fitness flux bound is tight near stationarity, when both the accumulation rate and the fitness flux approach 0. Figure computed using the Wright-Fisher model as in Fig. 1. The population size is fixed at *N* = 100. For technical reasons, the expected fitness flux curves were computed using an equivalent Moran model, see SI Sec. S10 and Fig. S2.

In summary, the accumulation of information is upper bounded by the KL cost of control, which in turn corresponds to the population size times the variation in fitness. However, if selection changes not only the probabilities of the final states, but also the paths that lead there (because it is strong, because adaptation is maintained for a long time, or because adaptation is reversed by time-dependent selection), then the information accumulated is less than the total cost.

### 3.5 Comparison with the fitness flux bound

The fitness flux theorem [35] implies another upper bound on information accumulation rate, Δ*D*(*X^t^*) ≤ 2*N*〈*φ*〉_*t*_, where 〈*φ*〉_*t*_ is the expected fitness flux per generation. It is plotted in gray in Fig. 4. It differs from the cost of selection bound both quantitatively and in terms of interpretation.

Quantitatively, neither bound is tighter in general. In Fig. 4B, the cost of selection bound is tighter in early stages of adaptation, and the fitness flux bound is tighter in the late stages. This is consistent with the interpretation of fitness flux as the rate of ongoing adaptation, or the rate of ascent in the mean fitness landscape/seascape [35]. This rate is high in the early stages of adaptation, when the population is far from the fitness peak and tends to climb up quickly. Later, when the population approaches a stationary distribution, there is no more adaptation on average, and 2*N*〈*φ*〉)_*t*_ as well as Δ*D*(*X^t^*) vanish. Meanwhile, the cost of selection bound *kN*〈*C*〉_*t*_ is tighter in the earlier stages when most of the cost is spent on new adaptation, but it remains positive under stationarity, due to maintenance costs.

Technically, the fitness flux theorem was originally derived in [35] under the diffusion approximation, and requires an additional assumption that the neutral process is at a stationary distribution with detailed balance. We derive and discuss the technical aspects of the fitness flux bound in SI Sec. S10 and Fig. S2, S3.

## 4 Maintenance of information

In this section, we ask how much information can be maintained in the genome for a given cost of selection. A general bound analogous to Eq. (10) seems to be out of reach for now, but we can study how the information maintained depends on key evolutionary parameters. We start by analyzing the single locus, two allele system, and then proceed to systems with large numbers of loci.

### 4.1 Single locus: weak selection is most efficient

Fig. 5A shows the information, *D*(*X*) and *D*(*G*), maintained by the single locus, two allele system at the stationary state under various strengths of selection. Stronger selection maintains more information – up to 1 bit at the genotype level, and more on the population level. However, it comes with a higher cost of selection 〈*C*〉, Fig. 5B. Notably, the cost increases faster then the maintained information. As a result, the amount of information maintained per unit cost decreases with selection strength, Fig. 5C.

**Figure 5:**
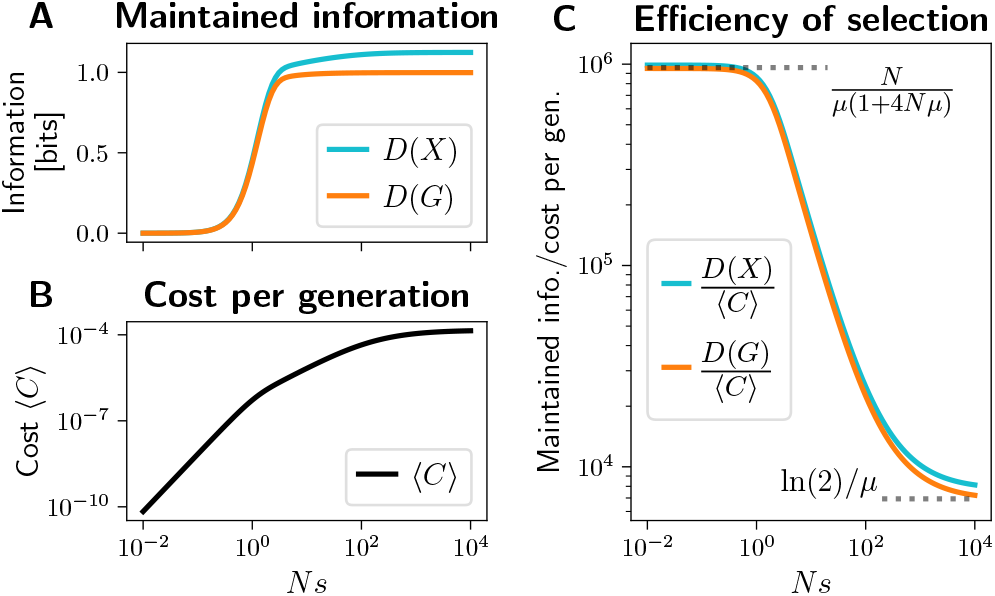
Maintenance of information in the single locus, two allele system. (A) The main plot shows the stationary values of information, *D*(*X*) (cyan) and *D*(*G*) (orange), as function of selection strength *Ns*. Stronger selection keeps the beneficial allele at higher frequencies, but this is associated with higher average cost of selection 〈*C*〉), shown in (B). Note that much of the time, one of the alleles is fixed and the cost C is zero. 〈*C*〉 is the average cost per generation over the stationary distribution of allele frequencies. (C) The ratio of the maintained information and the average cost of selection, *D*(*X*)/〈*C*〉 (cyan) and *D*(*G*)/〈*C*〉 (orange). Selection is most efficient when it is relatively weak (*Ns* ≪ 1), maintaining up to 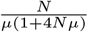 bits per unit cost at the genotype level, and inefficient when strong (*Ns* ≫ 1), maintaining only about ln(2)/*μ* bits per unit cost (dotted horizontal lines). The population size is *N* = 100 and mutation rate *μ* = 10^−4^.

There are two important asymptotic regimes. When selection is very strong, *Ns* ≫ 1, deleterious mutations are purged as soon as they arise, and *D*(*G*) ≈ 1 bit. Mutations arise with a probability *Nμ* per generation, and purging each costs *C* ≈ 1/(*N* ln(2)) (assuming truncation selection with *α* = 1 – 1/*N*, see SI Sec. S9). In this regime,

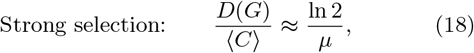

bits can be maintained per unit cost (see Fig. 5C). Similar arguments apply when *Nμ* > 1. The inverse scaling with *μ* is expected based on the deterministic mutation load [39] or Eigen’s error catastrophe [2] which occurs when selection cannot maintain sequences without error, and it was also derived by Watkins [5].

Selection is much more efficient when it is weak, *Ns* ≪ 1. Both the cost and the maintained information can be calculated under the diffusion approximation (see SI Sec. S4B for details). If mutation is also weak, *Nμ* ≪ 1, the amount of genetic variation (pairwise diversity) scales with 2*Nμ*, and the cost (variation in fitness) is approximately 〈*C*〉 ≈ *Nμs*^2^/(2 ln 2). Meanwhile, selection shifts the mean frequency of *A* away from 1/2 by about *Ns*/2, and this is associated with genotype-level information *D*(*G*) ≈ *N*^2^*s*^2^/(2 ln 2) bits. In this regime, up to *N*/*μ* bits per unit cost are maintained. When mutation *Nμ* is not negligible, a more accurate result is

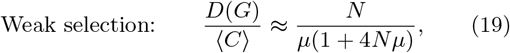

see SI Sec. S4. This limit is also highlighted in Fig. 5C. The special case when *Nμ* ≫ 1, 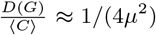 was previously derived by Watkins [5].

By itself, a single locus under weak selection cannot contribute much to biological function. However, selection can act on a polygenic trait influenced by many loci. If they are unlinked, we expect both the maintained information and the cost of selection to be approximately additive, and the ratio *D*(*G*)/〈*C*〉 to scale according to Eq. (19). To confirm this, we next study a polygenic system.

### 4.2 Information stored among many loci

We use an individual-based model to study a population of *N* haploids with *l* = 1000 biallelic loci, mutation and free recombination. Offspring are produced by sampling pairs of parents with selection, shuffling their genomes (at each locus, the allele from either parent is inherited with probability 1/2), and flipping each allele with probability *μ*. Selection acts on a fully heritable, additive trait with equal effects, *z_g_* = (the number of *A* alleles in *g*), with fitness being *w_g_* = (1 + *s*)^Z_g_^.

The results are shown in Fig. 6. The panel 6A shows an example of a stochastic population trajectory, indicating the phenotypes present in the population over time. The system is initialized with random genomes that contain the beneficial allele at each locus with probability 1/2, with *z* taking values around *l*/2 = 500 with binomial noise. Selection with *s* = 0.01 makes the beneficial alleles more frequent over time. The stationary distribution over phenotypes is in Fig. 6B. Under neutrality, *φ^Z^* = Binom(*l*, 1/2) by symmetry. The distribution *ψ^Z^* under selection is shifted relatively far from *φ^Z^*, leading to *D*(*Z*) = 88.0 bits of information on the phenotype level.

**Figure 6:**
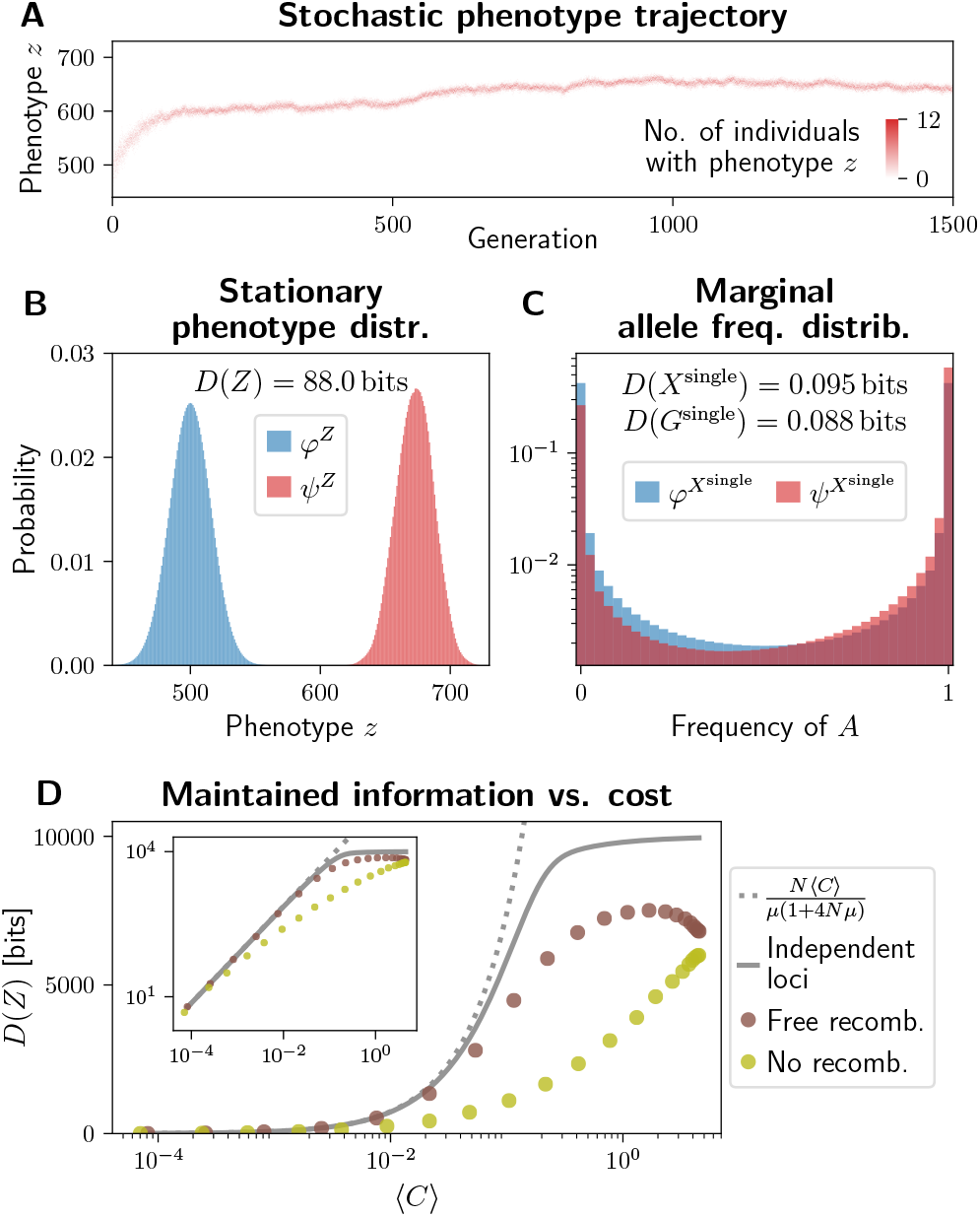
Maintenance of information in a system with *l* = 1000 biallelic loci. Selection is directional on an additive trait *Z* (= the number of beneficial alleles). (A) A heatmap showing the number of individuals in a population occupying each value of the phenotype *z* at each generation. The population is initialized as a collection of random genomes, each containing the beneficial allele at around *l*/2 = 500 loci. Over time, this number stochastically increases due to selection. Only the first 1500 generations of the trajectory are shown, the full trajectory was 5 × 10^3^ generations of burn-in and 2 × 10^5^ to estimate the stationary distributions in (B,C). (B) The stationary distribution over the phenotype *Z*, under neutrality (*φ^Z^*, blue) and selection (*ψ^Z^*, red), along with the phenotype-level information *D*(*Z*). Due to symmetry between loci and alleles, *φ^Z^*(*z*) = Binom(*z*; *l*, 0.5) is binomial. Under selection, *ψ^Z^* is obtained as the histogram over individuals and over 2 × 10^5^ generations at stationarity. (C) The marginal distribution over allele frequencies at individual loci, under neutrality (*φ*^*X*single^, blue, computed using a transition matrix for the single locus system) and under selection (*ψ*^Xsingle^, red, computed as a histogram over all loci and 2 × 10^5^ generations at stationarity). The associated *D*(*X*^single^) and *D*(*G*^single^) correspond to information maintained at one locus, and because the loci are approximately independent, the total information is about *l* = 1000 times more. The population size is *N* = 40, mutation strength *Nμ* = 0.02 and selection strength *Ns* = 0.4. (D) The relationship between the maintained information *D*(*Z*) and the cost of selection 〈*C*〉, with recombination (brown points), without recombination (olive points). This is compared with predictions under the assumption of independent loci (gray line; computed using single locus diffusion approximation and multiplying both information and cost by the number of loci) and the linear scaling with 〈*C*〉 based on Eq. (19) (dotted gray). Computed for a system with *l* = 10^4^ loci, population size *N* = 40, mutation strength *Nμ* = 0.02 and variable *Ns*. Distributions estimated from a stochastic trajectory over 5 × 10^4^ generations, after 5 × 10^3^ generations of burn-in. The inset shows identical data with a log vertical scale.

The population state distribution and the genotype distribution are inaccessible due to their dimensionality (see Fig. S1). However, we know that they are lower bounded by *D*(*Z*), which is easy to compute, and *D*(*Z*) ≈ *D*(*G*) since *Z* is the only trait under selection. Since the loci are unlinked and have equal effects, the information *D*(*Z*) can be divided evenly among them. The marginal distribution over allele frequencies is only slightly different from neutrality (Fig. 6C), by about *D*(*X*^single^) = 0.095 bits in terms of allele frequency distribution and *D*(*G*^single^) = 0.088 in terms of allele probabilities. The 1000 loci, however, combine to produce a large shift in the phenotype distribution, *D*(*Z*) ≈ 1000D(*G*^single^).

This information is maintained at a very low cost of selection, 〈*C*〉 = 0.0012 bits per generation, or relative fitness variance 〈*V*〉 = 0.0017. This amounts to *D*(*Z*) / 〈*C*〉 = 7.1 × 10^4^ bits per unit cost, only a little below the single locus limit *N*/*μ*/(1 + 4*Nμ*) = 7.4 × 10^4^ under weak selection.

### 4.3 Interference between loci

In practice, the selection on different loci might interfere, and this can hinder the maintenance of information. The interaction may be due to Hill-Robertson interference, linkage, or epistasis.

In Fig. 6D we vary the selection coefficient *s* on individual alleles in a *l* = 10^4^ locus system, and plot the maintained *D*(*Z*) against the cost 〈*C*〉. We use the individual-based model to compute these with free recombination (as in 6A-C) and with zero recombination (offspring genotypes are identical to those of single parents, up to mutation). We compare the results with the weak selection scaling according to Eq. (19), and results for 10^4^ loci that evolve independently (cost and information are summed over 10^4^ single locus systems).

With free recombination, weak selection maintains about as much information as if the loci were independent (brown points and gray line in Fig. 6D, inset), approximately according to Eq. (19) (gray dotted line). However, when selection is strong (〈*C*〉 ≈ 0.1 or more), individual alleles experience additional fluctuations in frequency due to random associations with alleles at other loci in a finite population [51], [52], reducing the efficiency of selection. As a result, the freely recombining loci maintain less information than if they were independent. This is in addition to the fact that under strong selection, maintenance is more costly even for independent loci (full gray line departs from dotted gray line, Fig. 6D). Extremely strong selection, which removes potentially adaptive variation at other loci, maintains even less information than more moderate selection, and it makes recombination ineffective (brown points at high 〈*C*〉 in Fig. 6D).

Without recombination, less information is maintained at any given cost (olive points in Fig. 6D). In fact, Watkins [5] has shown that due to clonal interference, organisms with no recombination cannot maintain more than the order of ln(*N*)/*μ* bits of information even if the cost is unlimited, making Haldane’s and Eigen’s results [2], [39] pertinent to asexual populations.

The advantage of recombination has also been recognized in a similar context by MacKay [4] and Peck and Waxman [6], and relates to the evolution of sex and epistasis. Recombination is advantageous when facing unconditionally deleterious or beneficial alleles [43], but can be disadvantageous when adaptation depends on beneficial combinations of alleles [53]. However, it is not clear if any form of selection can maintain more information at a given cost than 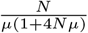 achieved by weak directional selection with recombination.

## 5 Discussion

Selection exerts control on evolving populations, but its capacity is limited. The limits to selection have been approached from various angles. Here we build upon previous work that had developed the idea that selection accumulates and maintains information in the genome [1], [2], and that this is associated with a cost in terms of variation in fitness, such as genetic load or fitness variance [39], [44]. The early work has suggested remarkably simple limits to selection: that the maximal rate of accumulation is bounded by the cost itself [1], [3], and that maintenance is limited to about 1/*μ* functional sites in the genome [2], [39].

Later work has pointed out that both accumulation [4], [7] and maintenance [5], [6] can exceed these limits, notably when recombination is involved. However, the general bounds remained unclear, possibly in part due to the difficulty of defining genetic information in general.

The measures of information that we have introduced in Sec. 2 coincide with or generalize previous definitions, and offer two advantages. First, they facilitate connections between different levels – e.g. between the abstract population-level information that has been studied theoretically in different contexts [34]–[36] and the effect that selection has on the distribution of phenotypes.

Second, the generality of our definition allows proving a general bound on information accumulation rate. This turns out to be a factor *N* faster than the early bounds, but depends on selection on individual loci being weak. The bound relies on a measure of cost of selection that connects the genetic load and fitness variance [48] with the KL cost in control theory [30], [31], recently used in the context of artificial selection [32].

How much information can be maintained in the genome at a given cost remains an open problem, but we have discussed how this might scale with the population size and the mutation rate. The scaling in Eq. (19) generalizes a result by Watkins [5] to realistic populations with *Nμ* < 1. Still, more work is needed to make claims about the information content of any real organism’s genome. Typical populations have *N_e_*/*μ* much greater than the genome size, suggesting that the genome size or other factors are more limiting than Eq. (19). The maintenance can be made more difficult by linkage or epistasis, and parts of the genome are likely under strong selection which is more costly. Still, Eq. (19) suggests that in theory, the genome could contain a substantial amounts of information among weakly selected loci, e.g. coding for polygenic traits. This is consistent with recent work [54] pointing out that mutation load does not pose severe limitations to the functional fraction of the human genome.

Similarly, the bound on accumulation rate in Eq. (10) hypothetically allows accumulation of information amounting to 10% of the human genome in about 10^6^ generations (6 × 10^8^ bits, assuming effective population size *N_e_* ≈ 10^4^, *k* = 2 and meager cost 〈*C*〉 ≈ 0.03 or relative fitness variance 〈*V*〉 ≈ 0.018 devoted to accumulation). But this is unlikely to have happened. Some selection was likely strong and more costly, and selection could have fluctuated, reversing previous adaptation. However, under the right conditions, information can accumulate very fast.

Our findings are complementary to the point raised by Kondrashov [41], that the survival of populations could be threatened by large numbers of weakly deleterious mutations (*Ns* < 1). While selection cannot purge them, it can perturb the allele frequency distribution of each by a small amount, and thus shift the distribution of higher level traits very far from neutrality. This is similar to the resolution by Charlesworth [55]. In fact, information accumulation and maintenance are most cost-efficient in this regime. This does not mean that a genomic architecture, where most mutations operate at *Ns* < 1 and information is encoded among many weakly specified sites, would evolve as an adaptation to maximise information gain. Nevertheless, such an architecture might arise in multicellular organisms as a side effect of their small effective population sizes and long genomes [56], [57].

Focus on the information content of genomes, rather than their fraction under selection, could help better frame the controversy sparked by some publications from the ENCODE project [12]–[16], [54], [58]. On the one hand, genomic regions under detectable selection (less than 15% in humans [59]) likely contain less than 2 bits per base pair, because their current function could be achieved by a number of alternative sequences (e.g. due to synonymous mutations in coding regions, or flexibility of transcription factor binding site sequence and location). On the other hand, regions without detectable selection could contain a considerable amount of information in the aggregate, at a low cost, encoding polygenic traits.

In bioinformatics, there already is a measure of information content applicable to short regulatory motifs [18], [19]. Future work could examine the precise relationship between this measure and our theoretical definitions. The generality of our framework also opens new directions for future research. One is to predict the maximal amount of information that can be maintained in genomes and populations with realistic parameters. Another is to study the information content of genomic elements with well-described genotypephenotype maps (e.g. promoters [26], [27]), under different hypotheses about selection on the phenotype.

## Acknowledgements

We thank Ksenia Khudiakova, Wiktor Młynarski and Sean Stankowski for discussions. G.T. and M.H. acknowledge funding from the Human Frontier Science Program (HFSP RGP0032/2018). N.B. acknowledges funding from ERC grant 250152 “Information and Evolution”.

## Supplementary information

### S1 Joint and conditional KL divergence, chain rule

For a single variable *U*, the KL divergence [1] between its distributions with and without selection is

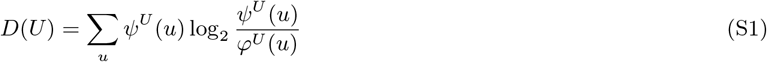

where *U* takes values *u* with probabilities *ψ^U^* (*u*) under selection and *φ^U^* (*u*) under neutrality.

To be well defined, the KL divergence requires that the support of *ψ^U^* is a subset of the support of *φ^U^*. In other words, if for some *u* we have *φ^U^*(*u*) = 0, then also *ψ^U^*(*u*) = 0 – outcomes impossible under neutrality are also impossible under selection. This condition needs to be respected when setting the initial conditions (*ψ^U^* and *φ^u^* at time zero). Over time, selection increases or decreases the probability of population states, genotypes or phenotypes that arise by reproduction with mutation, a by finite factor proportional to fitness. But selection cannot create entirely new states. On the other hand, some genotypes that arise by mutation can have zero fitness, and therefore be impossible under selection (*ψ^U^*(*u*) > 0 but *ψ^U^*(*u*) = 0). When this happens, the corresponding term *ψ^U^*(*u*) 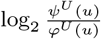 is set to zero. Therefore this is a very natural assumption, and analogous arguments apply to joint/conditional distributions which we discuss next.

For a pair of variables *U, V* we can write their joint and conditional KL divergence [1],

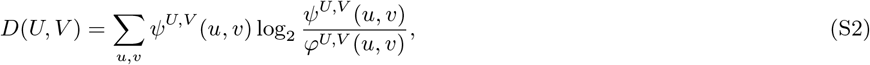

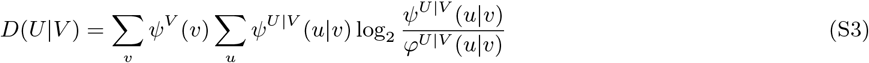

where *ψ^U,V^*(*u,v*) and *ψ*^*U*|*V*^(*u*|*v*) are the joint and conditional probabilities under selection, and *φ*^*U,V*^(*u,v*) and *φ*^*U*|*V*^(*u*|*v*) under neutrality. With these definitions, the chain rule states two possible decompositions of the joint KL divergence,

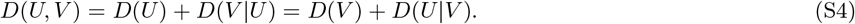

### S2 Population-level information: allele frequencies and LD

When there is a fixed number of loci, instead of genotype frequencies, an alternative way to describe a population state is in terms of allele frequencies. Allele frequencies by themselves, however, do not capture correlations between loci and therefore can miss some of the information that selection can accumulate on the population level. This can be expressed using the chain rule,

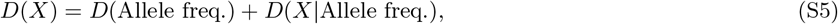

where the term *D* (Allele freq.|*X*) = 0 because regardless of selection, allele frequencies are fully determined by the genotype frequencies *X*. The term *D*(*X*|Allele freq.) quantifies how different from neutrality are the correlations between loci.

### S3 Violation of the bound by Worden 1995 by drift

The genotype-level information introduced by Worden [2] (see Eq. (11) there) is the KL divergence between the genotype frequencies and a uniform distribution,

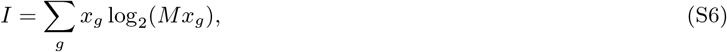

where *M* is the number of possible genotypes and *x_g_* the frequency of genotype *g* (denoted *q_j_* in [2]). Worden also introduced a similar genotype-level measure (Eq. (8) in [2], which was upper bounded by *I*. These measures of information can be seen as special cases of *D*(*G*) and *D*(*Z*) when there is no evolutionary stochasticity – *ψ^X^* is concentrated at a single value *x*, and *φ^X^* is concentrated at a single value of *x* that is uniform over all possible genotypes.

Worden proposes a bound on the rate of increase of *I* starting from a uniform *x*, and the maximal rate is proportional to a quantity similar to the genetic load, i.e. roughly a factor *N* (population size) times more stringent than the bound presented here.

The proof relies on the assumption that the population is large and *x*g evolve deterministically, but later, validity in finite populations is claimed (Sec. 2.6 in [2]). This is mistaken: in a realistic population, *I* can hardly be zero to start with, as there will be more possible genotypes than individuals and *x_g_* cannot be uniform. Starting from near uniform *x*, random drift will tend to remove variability from the population and concentrate all genotypes around some random ancestral genotype, and I will increase even without selection. This can also be seen as a consequence of the convexity of KL divergence: random fluctuations in *x* will, on average, increase *I*. This highlights the need to consider stochasticity as well as population variation when quantifying the intuitive notion of genetic information.

Worden’s stringent bound does hold if the genotype frequencies evolve deterministically and there is no recombination. This is consistent with our observation that selection accumulates information less cost-efficiently when *Ns* ≫ 1 and the fixation probability of a beneficial mutation is close to 1 (Main Text Fig. 3C,D).

### S4 The single locus, two allele system used for figures

We use a haploid single locus, two allele system to produce Figures 1-5. The figures are produced with a Wright-Fisher model, Moran model (only the fitness flux in Fig. 4, S2B and S3), and some intuition can be gained by approximating it as diffusion under weak selection. Note that this is only an illustration, more general classes of models are discussed in sections S5, S6 and S7.

The system has two alleles, *a* and *A*, where the latter is beneficial under selection. It is parametrized by the population size *N*, mutation rate *μ* and selection coefficient *s*(*x*) (which is frequency dependent only in Fig. 3C,D and S4).

#### S4A Wright-Fisher model

Under the Wright-Fisher model, the state space is a set of discrete frequencies of the *A* allele, *x_A_* = 0, 1/*N*, 2/*N*, …, 1, while the *a* allele always has the complementary frequency *x_a_* = 1 – *x_A_*. The two alleles have the following properties.

| Allele $g$ | frequency $x_g$ | fitness $w_g(x)$ | relative fitness $\hat{w}_g(x)$ |
| --- | --- | --- | --- |
| $a$ | $x_a = 1 - x_A$ | $w_a(x) = 1$ | $\hat{w}_a(x) = \frac{1}{1+s x_A}$ |
| $A$ | $x_A$ | $w_A(x) = 1 + s$ | $\hat{w}_A(x) = \frac{1+s}{1+s x_A}$ |

The probability of sampling allele *A* as a parent is *x_A_ŵ_A_*(*x*), and the probability of sampling it as offspring is

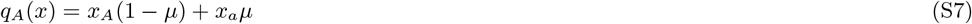

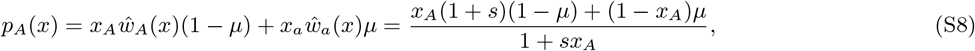

under neutrality and under selection respectively. The Wright-Fisher transition probabilities are given by the binomial distribution,

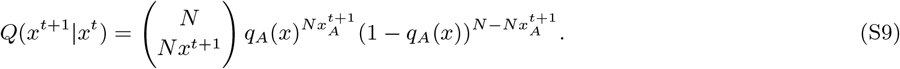

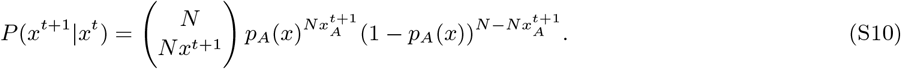

This is a case of the Wright-Fisher model with selection among parents (see Eq. (S27,S28) and SI Sec. S5A.2). An analogous discrete-time Moran model can be written by plugging Eq. (S7,S8) into Eq. (S43,S44).

All calculations were done with *N* ≤ 200. Given the small size of the system, we can compute the full matrix *P*(*x*^*t*+1^|*x*^*t*^), and calculate the distribution over genotype frequencies over time by iterating *ψ*^*Xt*+1^ (*x*^*t*+1^) = ∑_*x*_^*t*^ *ψ*^*Xt*^ (*x*^*t*^)*P*(*x*^*t*+1^|*x*^*t*^).

#### S4B The diffusion approximation

We write the diffusion approximation for the evolution of the frequency *x_A_*. Following the notation in SI Sec. S7, the first two moments of change of *x_A_* are given by

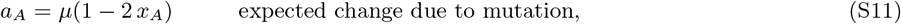

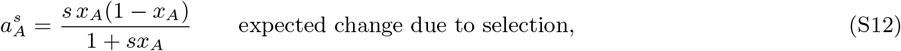

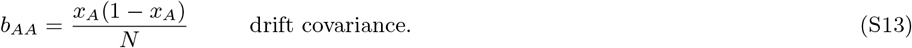

The diffusion equation for this system is a special case of Eq. (S57,S58).

##### Maintenance of information under weak selection

Since the diffusion process takes place along only one dimension, the stationary distribution can be determined by equating the probability flux to zero. For simplicity, we neglect the mean fitness 1 + *sx_A_* ≈ 1 in the denominator in Eq. (S12), by assuming that selection is weak, *s* ≪ 1.

The stationary distributions under selection and under neutrality are

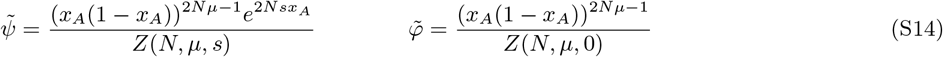

with the normalization constant

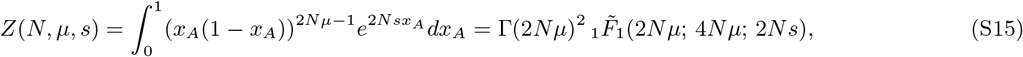

where Γ is the Gamma function and 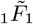 is the regularized confluent hypergeometric function.

Similar integrals yield results for the maintained information *D*(*X*), *D*(*G*) and the associated expected cost at the stationary state. We calculate the expectation of the cost 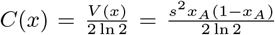 (see SI Sec. S9 and S7A). For the genotype-level information *D*(*G*), we also need the expected frequency of *A* which is equal to its marginal probability, 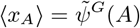. For brevity, we only write the leading terms in s for each quantity.

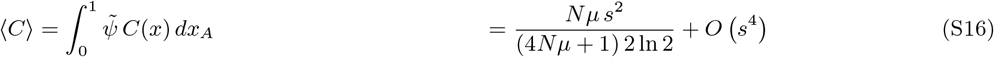

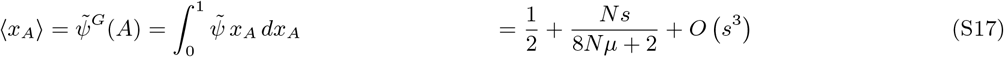

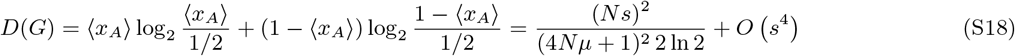

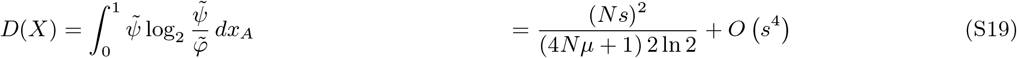

Notably, at weak selection, both the cost 〈*C*〉 and the information *D*(*X*), *D*(*G*) scale with *s*^2^. Their ratio is therefore given by the population size and the mutation rate,

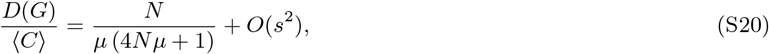

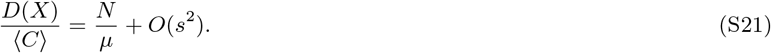

The ratio 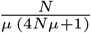 is shown in Fig. 5C.

### S5 The bound on information accumulation rate – Markov chains

The bound on information accumulation rate, as stated in Main Text Eq. (10,11) holds across several different model classes. Here we derive it for models that are Markov chain, in particular the Wright-Fisher model and the discrete Moran model. The two following sections contain similar derivations for continuous time Markov chains and the diffusion approximation. Note that all of the model parameters, such as those that describe selection, mutation or population size, can be time dependent, but we do not write it explicitly as we only need to focus on a single time step.

In the Markov chains class of models, the population state *X^t^* takes discrete values *x^*t*^* at discrete time steps *t*. The distribution over states is governed by

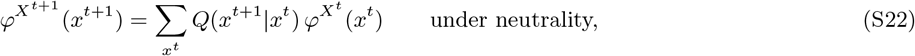

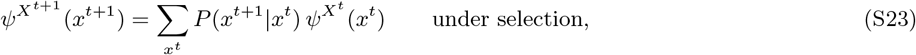

where *φ^X^* (*x^t^*) and *ψ^X^* (*x^t^*) are the marginal probabilities of population states at time *t*, and *Q*(*x*^*t*+1^|*x^t^*) = *φ*^*Xt*+ 1^|*X^t^* (*x*^*t*+1^|*x^t^*) and *P*(*x*^*t*+1^|*x^t^*) = *ψ*^*Xt*+ 1|*X*^ (*x*^*t*+1^|*x^t^*) are the transition probabilities. *Q*(*x*^*t*+1^|*x^t^*) and *P*(*x*^*t*+1^ |*x^t^*), as well as all the parameters that we later introduce to specify them, can be time-dependent, but we do not write it explicitly. The population-level information at time *t* is

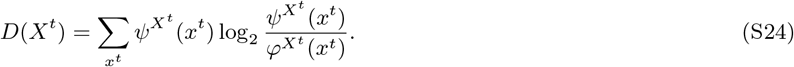

In general, the chain rule Eq. (S4) yields a bound

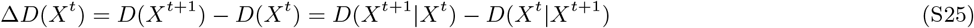

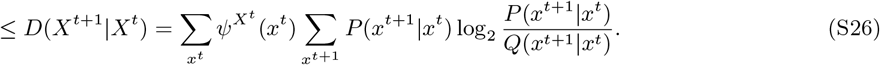

The expression ≤ *D*(*X*^*t*+1^|*X^t^*) corresponds to the expected KL cost of control [3], [4]. In the special case when *Q*(*x*^*t*+1^|*x^t^*) and *P*(*x*^*t*+1^|*x^t^*) are independent of time *ψ^Xt^* (*x^t^*) is the stationary distribution of *P*(*x*^*t*+1^|*x^t^*), *D*(*X*^*t*+1^|*X^t^*) is also the KL divergence rate between *P*(*x*^*t*+1^|*x^t^*) and *Q*(*x*^*t*+1^|*x^t^*). We now examine specific forms of the transition probabilities given by the Wright-Fisher model and the Moran model.

#### S5A Wright-Fisher model

In this general model, each time step *t* represents a generation, and consists of sampling a new population of *N* offspring genotypes that constitute the population at *t* + 1. The basic assumption is that the offspring genotypes are sampled independently with with probabilities *q_g_* (*x^t^*) without selection or *p_g_* (*x^t^*) with selection, leading to multinomial probability distributions over the frequencies *x*^*t*+1^ in the next generation,

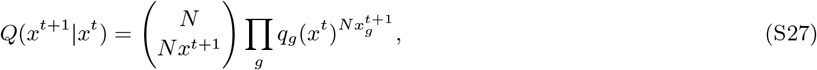

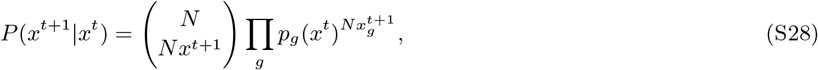

where 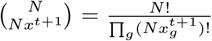 the multinomial coefficient. We can use these expressions to write down the general bound Eq. S26) as

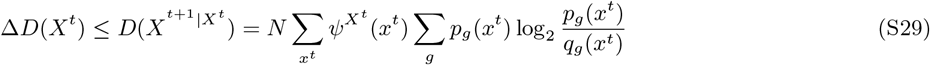

The probabilities *q_g_* (*x^t^*) capture arbitrary mutation and recombination, and we show examples of the form q_g_ (*x^t^*) can take below. Selection is modelled by the relationship between *q_g_* (*x^t^*) and *p_g_* (*x^t^*), and this can be done in two ways, which we discuss in the following subsections.

##### Asexual reproduction

In an asexual population with mutation, *q_g_*(*x*) will have the form

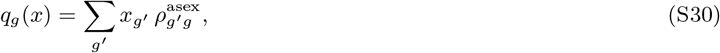

where 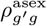 is the probability that a parent with genotype *g*′ produces offspring with genotype *g*, and includes arbitrary mutation.

##### Sexual reproduction, random mating

Provided that recombination happens always between two parental genotypes, we sum over possible pairs of parental genotypes,

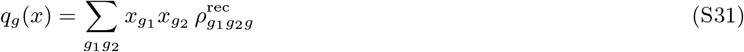

with *x*_*g*1_*x*_*g*2_ being the probability of a parental pair *g*_1_, *g*_2_ and 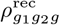 the probability that this pair produces offspring with genotype *g*. This includes arbitrary mutation and recombination.

Alternatively, we can distinguish between two sexes, classifying each genotype as male 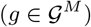 of female 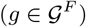. We sum over all male-female pairs,

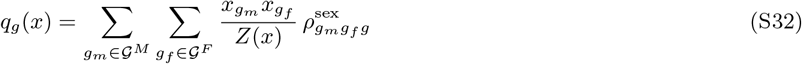

with 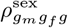 being the probability that parents *g_m_*,*g_f_* give rise to offspring *g* and

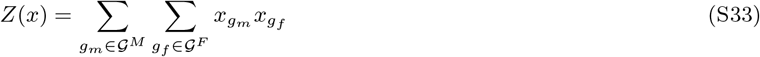

is a normalization factor such that *x_g_m__x_g_f__*/*Z*(*x*) is the probability of a parental pair *g_m_*,*g__f__*.

##### Nonrandom mating

In the case of sexual reproduction, we can replace the expression *x_g_m__*x*_g_f__*/*Z*(*x*) by a different expression for the probability of sampling a mating pair *g_m_, g_f_*. For example, we can include a factor 0 ≤ *σ_g_m_g_f__* ≤ 1 corresponding to the probability that individuals with this pair of genotypes will mate. Then *q_g_* (*x*) will have the form

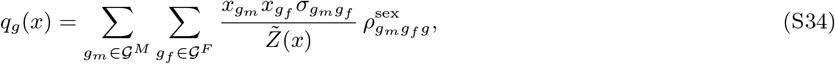

with normalization

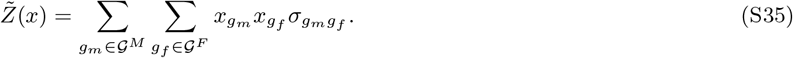

###### S5A.1 Selection in an infinite offspring pool

Here we assume that all individuals (under asexual reproduction) or all pairs of individuals (under sexual reproduction) in the population at time *t* contribute a large number of offspring to a common pool. The next generation then consists of *N* individuals sampled from this pool, with or without selection.

The genotype frequencies in the pool will be equal to *q_g_* (*x^t^*), since this is the probability that a random invididual (or pair of individuals) from the population *x^t^* has offspring with genotype *g*. Under neutrality, the genotypes that survive and constitute the next generation are sampled at random. Under selection, genotypes from the pool are sampled with probabilities proportional to fitness *w_g_* (*x^t^*), leading to

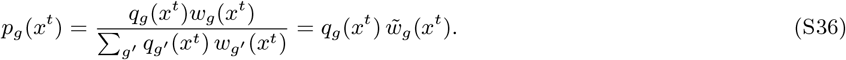

where 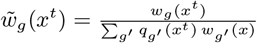 is the relative fitness of *g* calculated within the offspring pool where selection takes place.

Combining it with Eq. (S29), we obtain the bound

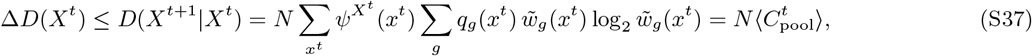

where the last expression coincides with the definition of the cost of selection in Main Text Eq. (S89), calculated at time *t* within the offspring pool where selection takes place, and averaged over possible population states *x^t^*.

###### S5A.2 Selection among parents

Here we assume that selection takes place before reproduction. From the population at time *t*, we first sample *N* genotypes (under asexual reproduction) or *N* pairs of genotypes (under sexual reproduction) as parents. These are sampled independently, with probabilities proportional to fitness. Then we sample one offspring genotype for each parent/pair of parents, with mutation and recombination, and these constitute the population at time *t* +1.

When sampling genotypes as parents, *g* gets picked with probability given by its frequency 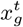 under neutrality, and 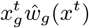 under selection where 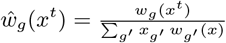, is the relative fitness of genotype *g*, now computed within the adult population at time *t*.

We can rewrite Eq. (S29) with

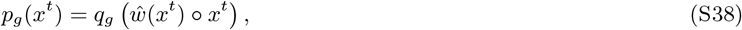

where 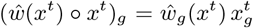 is the vector of genotype frequencies that are weighted by their relative fitness. However, Eq. (S38) is difficult to analyse. An easier approach is to introduce an intermediate variable *Y^t^*, which signifies the genotype frequencies among parents (or pairs of parents) at generation *t*. The Wright-Fisher model is a Markov chain of the form

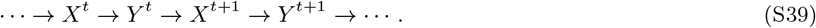

Selection operates in the step *X^t^* → *Y^t^* when parents are sampled from the existing population, but not in the step *Y^t^* → *X*^*t*+1^ where reproduction takes place with mutation and recombination. This can be expressed by writing the chain rule

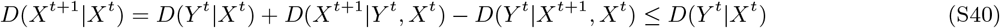

where the term *D*(*X*^*t*+1^|*Y^t^,X^t^*) = *D*(*X*^*t*+1^|*Y^t^*) = 0 because *Y^t^* → *X*^*t*+1^ is the reproduction step with no selection. The term *D*(*Y*^t^|*X*^*t*+1^,*X^t^*) is nonnegative and can be dropped since an upper bound on *D*(*X*^*t*+1^|*X^t^*) is sufficient for our purposes, but deserves some attention as it could make the bound loose. *D*(*Y^t^*|*X*^*t+1,Xt*^) can only be large when, given the genotype frequencies *X^t^,X*^*t*+1^ at two subsequent generations, there is uncertainty about the genotype frequencies among the parents *Y^t^* sampled from *X^t^* that gave rise to *X*^*t*+1^. In diverse populations with low mutation rates or large genomes, parents tend to be easy to identify, and *D*(*Y^t^*|*X*^*t*+1^,*X^t^*) will be small.

Finally, to calculate *D*(*Y^t^*|*X^t^*) we note that conditionally on *X^t^, Y^t^* is a multinomial variable with *kN* trials (*k* = 1 under asexual reproduction or *k* = 2 under sexual reproduction) and probabilities 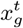 under neutrality or 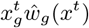 under selection. Then the bound can be written as

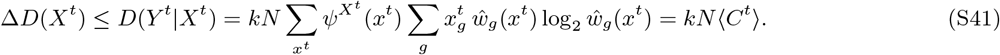

Here we have assumed no distinction between sexes, but we can extend to that case by sampling *N* parents of each sex separately and find

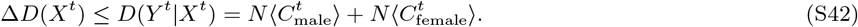

#### S5B Discrete-time Moran model

This model can be defined similarly to the Wright-Fisher model, but under the Moran model each time step consists of only one birth and one death, and there are *N* such time steps per generation.

The genotype that dies is chosen at random from the population *x^t^*, and the probability that it will be *g* is equal to its frequency 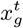. The probability that the genotype born is *g*′ is *q*_*g*′_(*x^t^*) under neutrality and *p*_*g*′_ (*x_t_*) under selection. These can take the same form as under the Wright-Fisher model, with selection within an infinite offspring pool or among parents. This gives rise to the transition probabilities

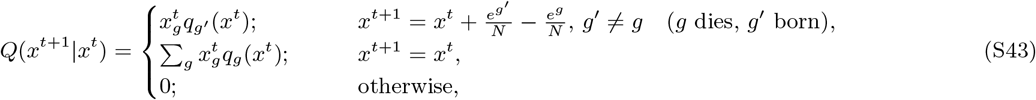

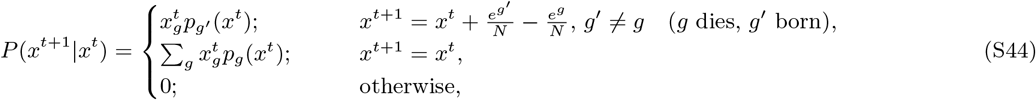

where*e^g^* is a vector of genotype frequencies with one at element *g* and zeros elsewhere. Now the bound on information accumulation rate, Eq. (S26), simplifies to

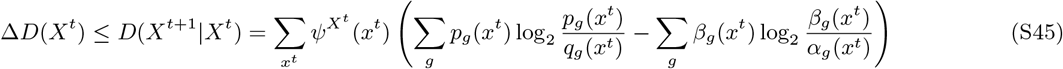

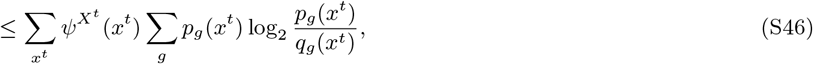

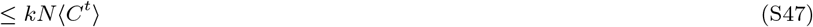

where we have used 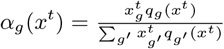, and 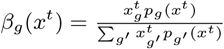 to denote the probability that if the genotype that is born and dies is the same, it is the genotype *g*. Even though selection makes *β_g_*(*x^t^*) different from *α_g_*(*x^t^*), such replacements leave the population unchanged regardless of g, and this is associated with the nonpositive second term inside the brackets in Eq. (S45), which reduces the amount of information that can be accumulated.

Eq. (S46) is almost identical to the bound on information accumulation rate under the Wright-Fisher model, Eq. (S29). This means that the discussion of the two models of selection in Sec. S5A.1 and S5A.2 also applies to the Moran model, which allows us to write Eq. (S47). The only difference is that in each time step, we sample only one genotype from the offspring pool, or one parent (or pair of parents). This is reflected in the missing factor *N*. As there are *N* time steps per generation, the bound on information accumulated per generation again scales with *kN*〈*C*〉^*t*^).

Note, however, that the effective population size of the Moran model (i.e. the population size of a Wright-Fisher model with the same covariance in allele frequency change per generation) is N_e_ = N/2. This is because in the Moran model, both the births and deaths are random events, whereas the Wright-Fisher model only has random births. We can therefore expect the tighter bound Δ*D*(*X^t^*) ≲ *kcN_e_*〈*C^t^*〉) to hold for large enough populations, as both models approach the same diffusion limit. We also prove the bound under the diffusion approximation separately in Sec. S7.

### S6 The bound on information accumulation rate – continuous-time Markov chains

In this class of models, the genotype frequencies *X^t^* are discrete, but the time *t* is continuous. The distributions *φ^X^* (*x*), *ψ^X^* (*x*) over *X^t^* are governed by the master equations

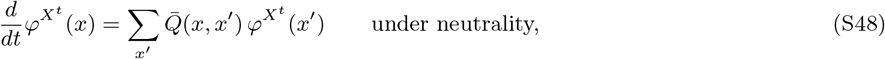

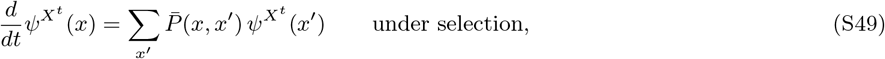

where 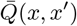 and 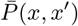 are transition rates from *x*′ to *x* under neutrality and selection respectively. These can be time dependent. The population-level information *D*(*X^t^*) is defined as in Eq. (S24), but now changes continuously in time. The rate of this change is upper bounded as

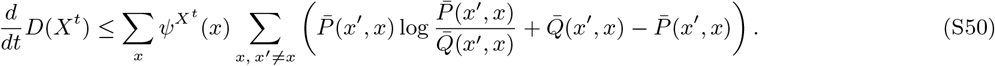

When 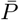 and 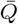 are independent of time and *ψ^X^* (*x*) is the stationary distribution associated with 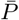, the right hand side corresponds to the KL divergence rate between 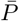 and 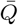, as derived in [5]. The bound Eq. (S50) can be verified algebraically, or derived from the discrete bound Eq. (S26) by taking 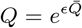, 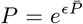 and the limit *ϵ* → 0. We now show an example of the form that 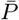 and 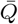 can take.

#### S6A Continuous-time Moran model

This model is based on its discrete-time counterpart in Sec. S5B. Only transitions consisting of replacing one genotype (*g*, death) by another (*g*′, birth) are allowed, and the transition rates have the form

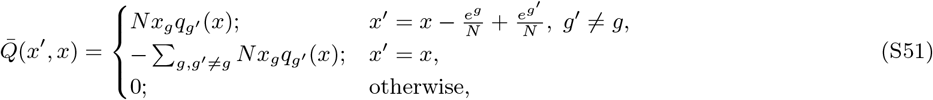

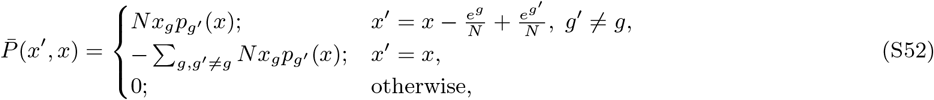

where time is measured in generations, i.e. there are on average *N* replacement events per unit time. With this form of the transition rates, we can rewrite the general bound Eq. (S50) as

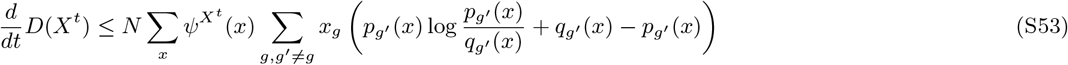

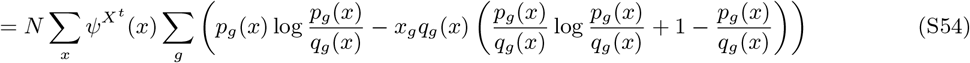

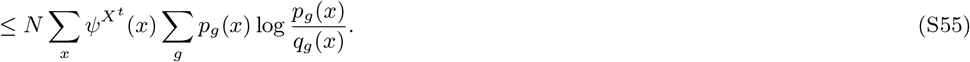

This is the same bound as for the discrete Moran model in Eq. (S47), up to the factor *N*, which is due to the different unit of time.

### S7 The bound on information accumulation rate – diffusion approximation

Here we show an upper bound on the rate of accumulation of information under the diffusion approximation. The approach is similar to Iwasa [6] (who assumed detailed balance) and Hasegawa [7] (who did not), but here we distinguish between the processes with and without selection. We start by deriving a general bound for a pair of diffusion processes, and then apply it to the population genetics context in Sec. S7A.

For brevity, we write the probability density over population states *x* at time *t* as *φ* = *φ*(*x,t*) under neutrality and *ψ* = *ψ*(*x,t*) under selection. The population-level information is now determined by integration,

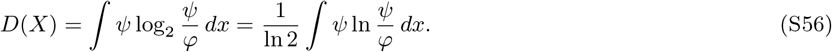

Note that while we stick to measuring information in bits, it is more convenient to use the natural logarithm during the derivation.

The diffusion equation is parametrized by the first and second moment of change in *x_g_*. Selection is assumed to only exert control through the first moment, which we label as *a_g_* under neutrality and 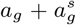 under selection. The second moment is *b*_*gg*′_, both under neutrality and under selection. All these are functions of *x* and *t*, e.g. *a_g_* = *a_g_*(*x,t*), but we do not write this for brevity. We sum over any index that appears twice in a term, e.g. 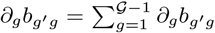. Note that the diffusion is described in the subspace of 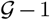 genotype frequencies, where 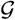 is the number of genotypes – the last frequency is determined by normalization, 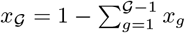. The diffusion equation is

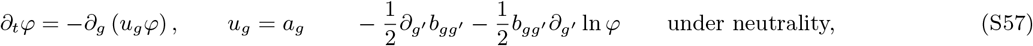

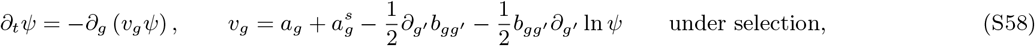

where we introduced the velocity fields *u_g_* = *u_g_*(*x,t*) and *v_g_* = *v_g_*(*x,t*) such that *u_g_φ* and *v_g_ψ* are the probability fluxes under neutrality and under selection respectively. From their definition, it follows that

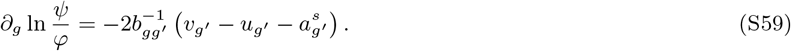

The rate of change of *D*(*X*) can be written as

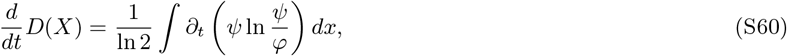

and the integrand can be written as

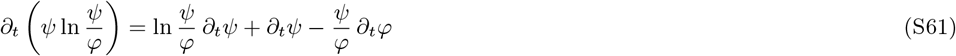

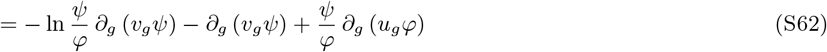

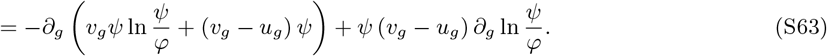

The first term is a divergence and vanishes after integration in Eq. (S60) because *u_g_φ* and *v_g_ψ* cannot cross the domain boundary (assuming *ψ/φ* < ∞). Therefore

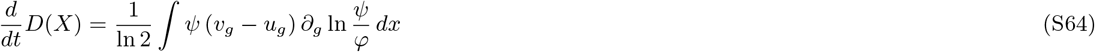

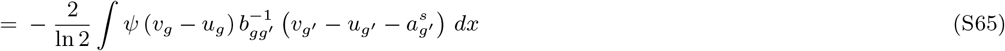

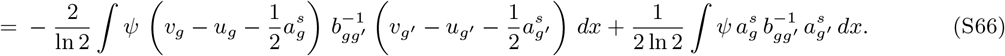

The last expression has two terms, both of which are quadratic forms. The first one makes a nonpositive contribution, leading to the upper bound in information accumulation rate,

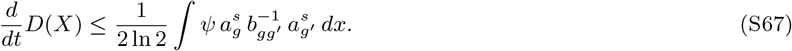

On the right hand side we can identify the KL cost of control [4] in bits. This bound holds for any pair of diffusion processes with the same fluctuations covariance *b*_*gg*′_. We will now discuss it in the context of population genetics.

#### S7A Application to population genetics

The bound Eq. (S67) does not depend on the form of *a_g_*, and therefore it can be used to model arbitrary mutation and recombination, for example by taking ation and

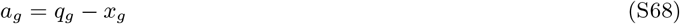

with *q_g_* = *q_g_* (*x*) as introduced in the discrete models above. Notably, *a_g_* does not need to be the gradient of a scalar potential.

The fluctuations in genotype frequencies can modeled based on multinomial sampling in the Wright-Fisher model (Eq. (S27,S28)),

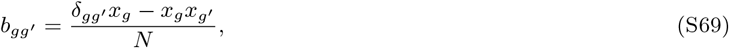

with no summation over *g*, and where the (effective) population size *N* can be time-dependent. This has the inverse [8]

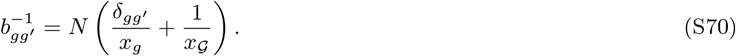

The control term 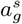 imposed by selection can be written as

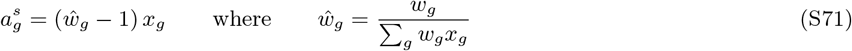

where *w_g_* is the fitness of genotype *g*, possibly time and frequency dependent, and *ŵ_g_* is the relative fitness. With these definitions, we find that

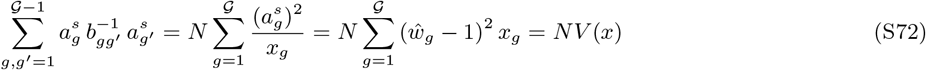

and the bound on information accumulation rate is

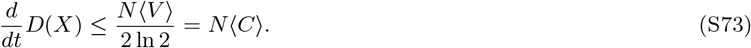

In the last equation, we equate 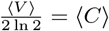 – we show that this holds under weak selection in Sec. S9.

We can get intuition about the tightness of this bound by analyzing the first, nonpositive term in Eq. (S66). The bound is only tight when 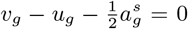 for all *x* with nonzero *ψ*. An interesting specific case is when the neutral process is at an equilibrium with detailed balance, such that *u_g_* = 0. We note that *v_g_* can be decomposed as 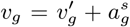 into the contribution from selection 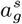 and all other evolutionary forces, 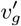. Our bound is then tight when 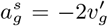, i.e. when selection induces a probability flux in exactly the opposite direction and exactly twice the magnitude as the all the other evolutionary forces combined. While this might occasionally and approximately be the case, it will only be a transient phenomenon.

Suppose that the population starts at the neutral equilibrium with 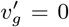 and then selection starts to act, e.g. after a change in the environment. For any nonzero 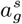, the bound cannot be tight as 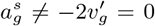. After some time a new equilibrium might be reached, with 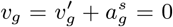, where again, the bound is not tight 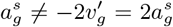. In this case, maintenance costs are incurred but no further adaptation takes place. The bound can only be tight for a moment when adaptation is taking place, selection pulls the population in the opposite direction as the other evolutionary forces combined, but selection is twice as strong, 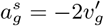.

### S8 Relationship with free fitness and statistical physics

Stochastic models in population genetics show some mathematical properties analogous to statistical physics. In particular, a quantity called free fitness, analogous to free energy in physics, can be defined and shown to monotonically increase over time. In this section we provide some background about free fitness and discuss two connections with our work. First, the increasing property of free fitness can be proved by a method similar to the proof of our bound on information accumulation rate. Second, free fitness can be written as the difference between mean log fitness and genetic information as defined in this paper (e.g. on the genotype or population level), implying that evolution tends to maximize mean log fitness at a given amount of information.

#### S8A Boltzmann form of stationary distributions

Under suitable conditions, the stationary distributions in population genetics models take a form similar to the Boltzmann distribution. Models on both the population level and the genotype level display this property.

- If mutation is weak (*NU* ≪ 1 where *U* is the total mutation rate across the studied genomic region), populations are mostly monomorphic, with only occasional fixations of a different genotype. The system can then be described with the most recently fixed genotype *g* and the distribution *ψ^G^*(*g*). The stationary distribution 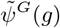 takes the form [9], [10]

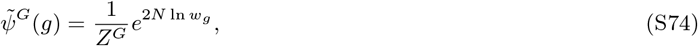

where 2*N* is again analogous to inverse temperature, log fitness ln *w_g_* is analogous to negative energy, and *Z^G^* is normalization constant.
- Assuming many biallelic loci under linkage equilibrium, we can describe the system with the vector of allele frequencies *p* and the joint distribution *ψ^P^* (*p*) over them. The stationary distribution 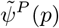 can be derived from the diffusion approximation and takes the form [11], [12]

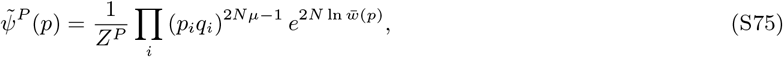

where *p_i_* and *q_i_* = 1 – *p_i_* are the allele frequencies of the two alleles at locus *i* and *Z^P^* is a normalization constant. Twice the population size 2*N* takes the role of inverse temperature and log mean fitness ln 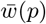 takes the role of negative energy. The factors (*p_i_*(1 – *p_i_*))^2*Nμ*–1^ correspond to mutation and drift potential (similar to e.g. chemical potential), which will be made clearer in the next subsection.

The formulas apply to haploids, but similar formulas apply to diploids or when mutation coefficients vary across loci or alleles (see e.g. the SI of [10]). Importantly, they depend on the assumption of detailed balance. At stationarity, net probability flux between any two allele frequency vectors or genotypes must be zero, making 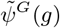 and 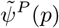 equilibrium distributions. This can be violated under certain forms of mutation, recombination and strong selection, leading to additional terms related to robustness [13].

#### S8B Free fitness

When a system starts from an arbitrary initial distribution and approaches the stationary distribution in Eq. (S75) or Eq. (S74), we can track this progress using free fitness [6], [10], which increases monotonically in time as was shown previously and as we can also prove in more generality here.

- On the genotype level, we can define free fitness *F^G^* at any distribution *ψ^G^*(*g*) away from equilibrium as a sum of expected log fitness and entropy terms,

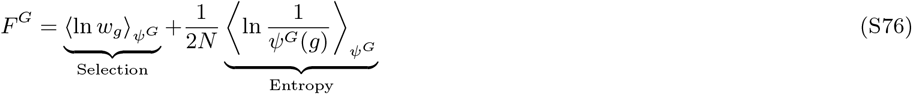

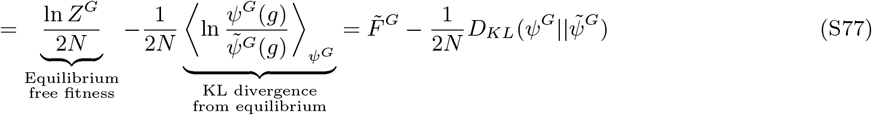

where 〈·〉_*ψ^G^*_ denotes an expectation over *g* ~ *ψ^G^*(*g*). In the special case of equilibrium 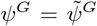, we obtain 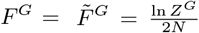, and away from equilibrium, free fitness is reduced by an amount proportional to the KL divergence between the actual distribution *ψ^G^* and the equilibrium 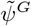.
- On the population level with linkage equilibrium, free fitness at a distribution *ψ^P^* can be defined as a sum of three terms – a negative potential for selection, mutation and drift, and entropy:

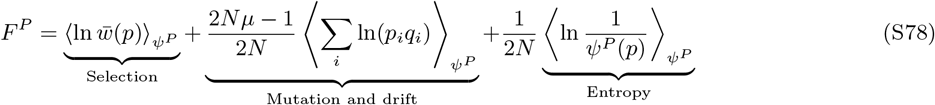

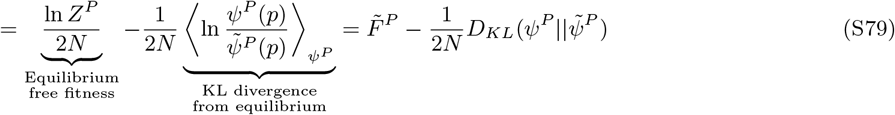

where the expectations 〈·〉 are taken over *p* ~ *ψ^P^*(*p*). While mutation and drift now appear as additional terms in free fitness, free fitness can again be decomposed into its value at equilibrium and a difference proportional to the KL divergence away from it.

The key property of the free fitness is that it is a non-decreasing function of time, until it is maximized at equilibrium. This was proved by Iwasa [6] for the case of *F^P^* and Sella and Hirsh [10] for the case of *F^G^*. In the next section we show that both results can also be derived by the same method as our bound on information accumulation rate.

#### S8C Monotonic convergence of stochastic processes to their stationary distributions

The general bounds on information accumulation rate (Eq. (S26) for Markov chains, Eq. (S50) for continuous time Markov chains and Eq. (S67) for the diffusion approximation) apply for any pair of stochastic processes, provided that they have compatible support such that the KL divergence is well defined. To make this explicit, we focus on the case of discrete Markov chains, consider some general process instead of the neutral process *φ*, and introduce notation that generalizes *D*(*X^t^*),

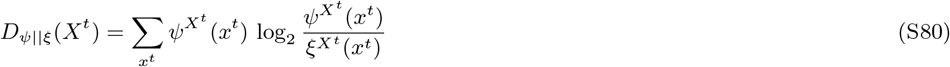

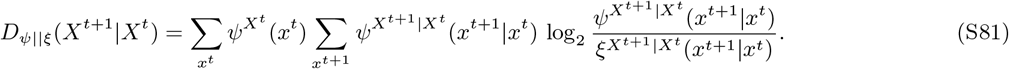

The KL divergence chain rule now yields the inequality

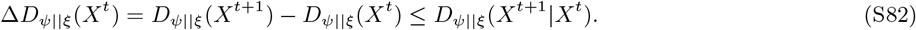

If *ξ* = *φ* is the neutral process, this is the KL cost of selection bound on the information accumulation rate (Eq. (S26) and Main Text Eq. (9). But we can also choose such that it does contain selection and has the same transition probabilities as *ψ*, but starts from a different initial condition, i.e. *ξ*^*Xt*+1|*Xt*^ = *ψ*^*Xt*+1|*Xt*^ and *ξ*^*X*0 ≠ *ψX*0^. Then we find that *D*_*ψ*||*ξ*_(*X*^*t*+1^|*X^t^*) = 0 and Δ*D*_*ψ*||*ξ*_(*X^t^*) ≤ 0, i.e. the divergence between *ψ*^*Xt*^ and *ξ*^*Xt*^ is non-increasing over time, because relative to *ξ*, there is no control exerted on *ψ*.

If, in addition, the system has a unique stationary distribution 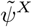 and *ξ*^*X0*^ is initialized there (and therefore stays there indefinitely, 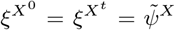 for any *t*), we find that *ψ*^*Xt*^ converges to this stationary distribution 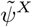 monotonically in terms of the KL divergence *D*_*ψ*||*ξ*_(*X^t^*). Similar proofs apply to continuous time Markov chains and diffusion, since we only need to replace *φ* by *ξ* and repeat the derivation leading to Eq. (S50) and Eq. (S67).

In the two regimes discussed above in Sec. S8A, the population state *X* corresponds to some fixed genotype *G* or a vector of allele frequencies *P*. Therefore 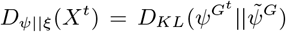 or 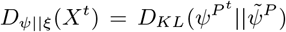 are non-increasing functions of time. Together with Eq. (S77,S79), this implies that *F^G^* or *F^P^* are non-decreasing functions of time.

Iwasa [6] and Sella and Hirsh [10] proved the same result by different methods. In our framework it emerges as a special case of the information accumulation bound with zero control. The key part of our proof, stating that *D*_*ψ*||*ξ*_(*X^t^*) is non-increasing, is also more general (regarding the state space, the form of the stationary distribution, and detailed balance – although free fitness is not defined so generally). A similarly general proof for continuous time Markov chains, as well as several related results for replicator dynamics and reaction networks, is reviewed in reference [14].

#### S8D Free fitness as a trade-off between fitness and information

We can rewrite the expressions for free fitness using the genotype and population-level information respectively. We first write down the neutral stationary distributions. On the genotype level, we assume that it is uniform over 4^*l*^ possible sequences of length *l*,On the population level,

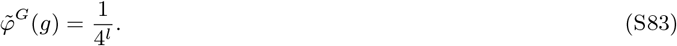

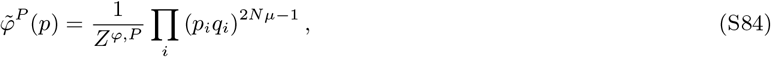

where *Z^Φ,P^* is the normalization constant, to be distinguished from *Z^P^* in Eq. (S75) which includes selection.

Using 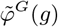 and 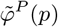, we can rewrite free fitness, Eq. (S77,S79), as

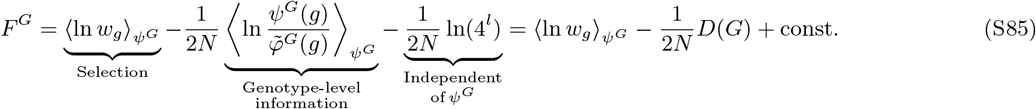

on the genotype level and

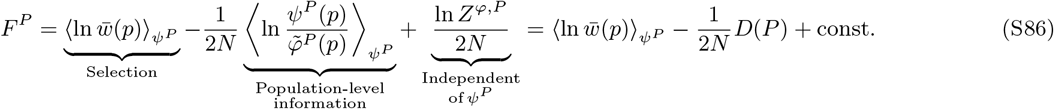

on the population level. In both cases we have emphasized that terms independent of *ψ^G^* or *ψ^P^* are constant in time and therefore not important for the dynamics of free fitness. Up to the constant, this formula for free fitness has also been used in the paper on fitness flux [15]. The fitness flux theorem (ref. [15] and Sec. S10) then provides perhaps the most elegant proof that free fitness is a non decreasing function of time, as it relates changes in *D*(*P*) to changes in expected fitness.

Free fitness tends to increase over time until it is maximized at the Boltzmann-like equilibrium distribution 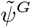 or 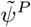. In other words, evolution maximizes the expected log fitness while constraining the amount of genetic information, with 1/(2*N*) serving as a Lagrange multiplier that controls the trade-off.

### S9 Properties of measures of cost of selection

Here we prove general inequalities between the genetic load *L*(*x*), relative fitness variance *V*(*x*), and the information theoretic cost *C*(*x*). We also derive the form of *C*(*x*) for the special cases of weak selection and truncation selection. The three measures are defined as

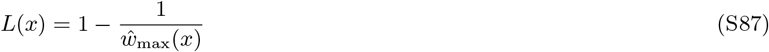

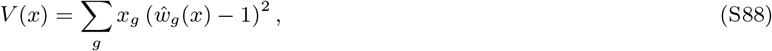

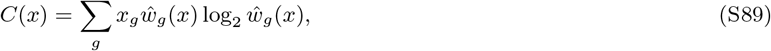

where *x_g_* is the frequency of genotype *g* in the population and *ŵ*_max_(*x*) = max_*g*; *x_g_ > 0*_ *ŵ_g_*(*x*) is the relative fitness of the fittest individual that is present in the population (*x_g_* > 0).

We note that some previous work has defined *ŵ*_max_(*x*) to be the maximum fitness possible, i.e. the fitness of an ideal genotype with no deleterious mutations regardless of whether such an individual exists. Load computed with such a definition is higher, and this has led to claims of severe restrictions on the rate of adaptive substitutions [16] and the functional fraction of the human genome [17]. However, load under this definition has been criticized as irrelevant, since the ideal genotype has a vanishing probability of existing in the population, and if only the fitness values likely to be present in the population are considered, load-based restrictions are more permissive [18], [19]. Our definitions of *L*(*x*), *V*(*x*) and *C*(*x*) all focus on the existing variation of fitness in the population *x*. *L*(*x*) is also related to the concept of lead, which was defined as the difference between the maximum and the mean log fitness in a traveling wave [20].

#### Truncation selection and limitations by reproductive capacity

Under truncation selection, a fraction *α* of individuals in the population has constant relative fitness, equal to the maximum *ŵ_g_* (*x*) = *ŵ*_max_(*x*) and the remaining fraction 1 – *α* has relative fitness zero *ŵ_g_* (*x*) = 0. By definition, the mean relative fitness must be ∑_*g*_ *x_g_ ŵ_g_* (*x*) = 1, which requires *ŵ*_max_(*x*) = 1/*α*. The three measures of cost of selection then are

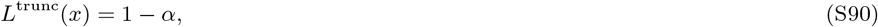

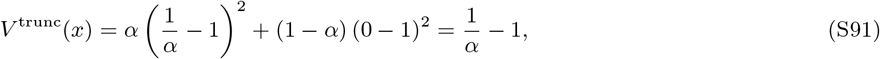

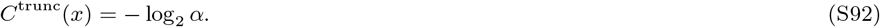

At a constant population size, the expected number of offspring of an individual is equal to their relative fitness *ŵ_g_* (*x*) (or 2*ŵ_g_* (*x*) under sexual reproduction, with two parents per offspring). In a species with a reproductive capacity *R*, we have *ŵ*_max_(*x*) ≤ *R* and the load is limited as *L* ≤ 1 – 1/*R*. *V* (*x*) and *C*(*x*) at given *R* are maximized under truncation selection, when only the most extreme relative fitness values available are occupied (*Nα* individuals have relative fitness *ŵ_g_* (*x*) = 1/*α* = *ŵ*_max_ (*x*) = *R* and *N* – *Nα* individuals have fitness 0). This implies upper bounds *V* (*x*) ≤ *R* – 1 and *C*(*x*) ≤ log_2_ *R*.

#### General inequality between *L*(*x*) and *V*(*x*)

From Eq. (S87), the genetic load *L*(*x*) determines the maximum relative fitness in the population, *ŵ*_max_(*x*) = 1/(1 – *L*(*x*)). Given that relative fitness of all individuals in the population must lie between 0 and *ŵ*_max_(*x*), its variance *V*(*x*) is maximized when only these extreme values are occupied, i.e. under truncation selection. In that case we have *V*^trunc^(*x*) = 1/*α* – 1 with *α* = 1/*ŵ*_max_(*x*). This implies a general bound,

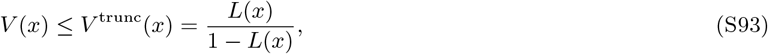

with equality under truncation selection. The same inequality was derived in ref. [21] by other means.

#### General inequality between *L*(*x*) and *C*(*x*)

Since logarithm is an increasing function and *x_g_ŵ_g_* (*x*) ≤ 0, we can upper bound each term of the form *x_g_ŵ_g_* (*x*) log_2_ *ŵ_g_* (*x*) in Eq. (S89) by *x_g_ŵ_g_* (*x*) log_2_ *ŵ*_max_(*x*). Summing over *g*, we obtain

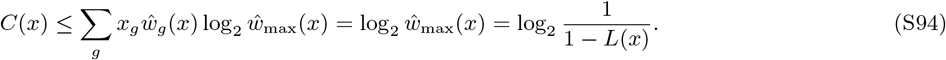

Equality is again achieved under truncation selection.

#### General inequality between *V*(*x*) and *C*(*x*)

We use the inequality 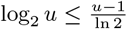 in Eq. (S89) to obtain

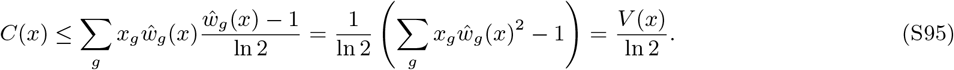

Equality is approached under truncation selection when *α* → 1.

#### *C*(*x*) under weak selection

Here we assume that for all genotypes *g* present in the population (*x_g_* > 0), the relative fitness *ŵ_g_* (*x*) is close to 1. We can then use the Taylor expansion

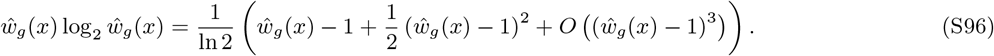

Combining this with Main Text Eq. (S89), we find

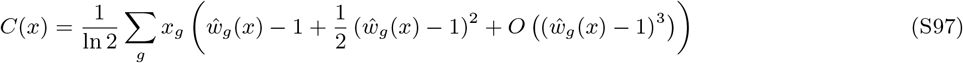

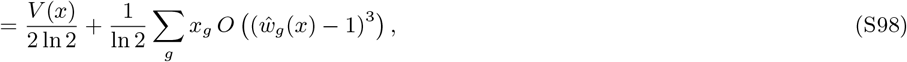

or in short, *C*(*x*) ≈ *V*(*x*)/(2 ln 2) under weak selection. This is particularly relevant for the diffusion limit, when the population size is sent to infinity, and the selection strength is rescaled inversely to the population size.

### S10 Fitness flux theorem

In this section we compare the newly introduced bound on information accumulation rate (the *cost of selection bound*) and a similar bound implied by the fitness flux theorem [15] (the *fitness flux bound*). The fitness flux theorem was originally derived under the diffusion approximation. For better comparison, we also derive an analogous result for discrete-time Markov chains. We then discuss the distinct interpretation of the two bounds, and illustrate them (using both the discrete and diffusion expressions) in Fig. S2.

#### S10A Discrete-time Markov chains

The fitness flux theorem, like its counterparts in statistical physics (e.g. [22]), is based on the comparison of forward and reverse path probabilities. For simplicity, we will not derive the fitness flux theorem in its general form, but rather the form that allows a direct comparison with the cost of selection bound.

We focus on short paths consisting of only one step, (*x_t_*, *x*^*t*+1^). The probability of the forward path (*x_t_*, *x*^*t*+1^) is *ψ*^*Xt*^ (*x_t_*)*P*(*x*^*t*+1^|*x_t_*). We consider a probability distribution over reverse paths, *ψ*^*Xt*+1^ (*x*^*t*+1^)*P*(*x^t^*|*x*^*t*+1^), which is normalized to 1 – we can write this as

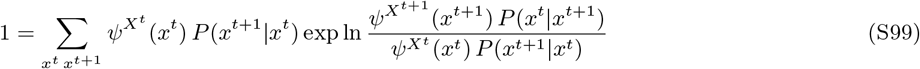

By Jensen’s inequality,

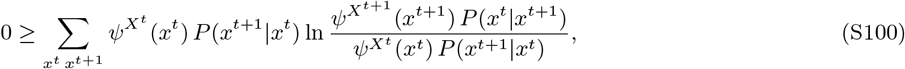

Next, inside the logarithm, we divide and multiply by the neutral probabilities,

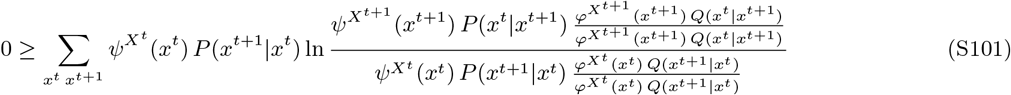

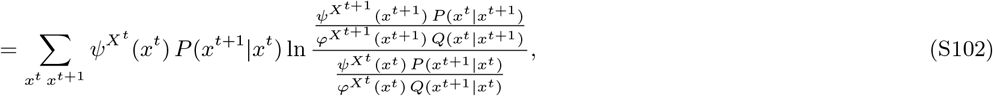

where we assumed that the neutral process is at a stationary distribution with detailed balance, i.e. *φ*^*Xt*^ (*x^t^*) *Q*(*x*^*t*+1^|*x^t^*) = *φ*^*Xt*+1^) *Q*(*x*|*x*^*t*+1^). Finally, we rearrange terms and divide by ln 2 to get an expression in bits,

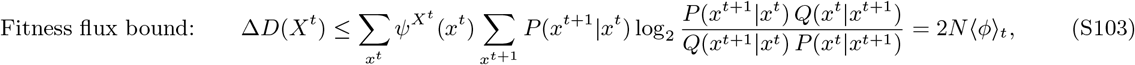

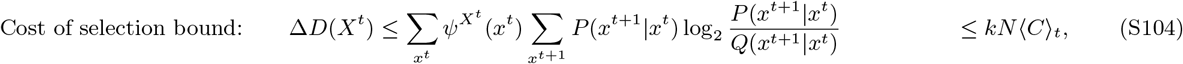

where is the 〈*ϕ*〉_*t*_ discrete analog of the fitness flux, averaged over the possible transitions (*x^t^*, *x*^*t*+1^),

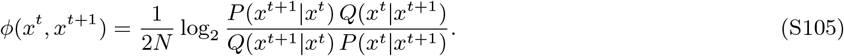

The interpretation of this expression and the relationship to fitness accumulation is most clear in the diffusion approximation, see below. To help compare the fitness flux bound with the cost of selection bound, we take the expectation over the final state *x*^*t*+1^ and compute the expected fitness flux from any initial state *x^t^*, *ϕ*(*x^t^*) = ∑_*x*^*t*+1^_ *P*(*x*^*t*+1^|*x^t^*)*ϕ*(*x^t^*, *x*^*t*+1^). An example plot of *ϕ*(*x^t^*) is in Fig. S2A. For comparison, we also included in Eq. (S104) the cost of selection bound.

While the cost of control is a non-negative conditional KL divergence, the fitness flux bound contains an additional term related to the reverse transition probabilities, and can be negative (this is more easily interpretable as the mutation term in the diffusion approximation). The fitness flux bound relies on the additional assumption that the neutral process is at a stationary distribution with detailed balance. This can be satisfied in the single locus, two allele system when using the Moran model, but it is violated by the Wright-Fisher model which we use throughout most of the paper.

#### S10B Diffusion approximation

Mustonen and Lässig [15] derive the fitness flux theorem using a similar method but under the diffusion approximation, where the fitness flux is related to the rate accumulation of fitness. We include here an informal account of how that relates to the formula in Eq. (S104).

In continuous time, we can generalize the definition of fitness flux in Eq. (S105) to an arbitrary time interval Δ*t* and the transition (*x^t^*, *x*^*t*+Δ*t*^).

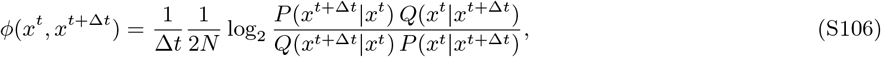

where we also divided by Δ*t* to get the fitness flux per generation. Under the diffusion approximation, if Δ*t* is small, the transition probabilities *P, Q* will be approximately normal with parameters given by *a*(*x^t^*), *a^s^*(*x^t^*) and *b*(*x^t^*) (see Sec. S7; we drop the dependence on *x^t^* for brevity),

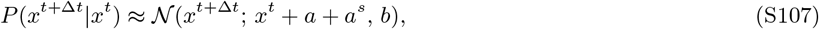

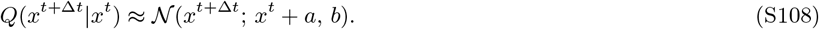

Then in Eq. (S106) we recover the definition of fitness flux from [15],

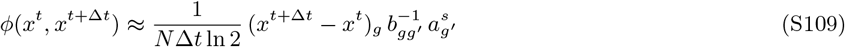

where we sum over repeated indices as in Sec. S7. Note that the factor 1/ln 2 appears because we use base 2 logarithms throughout. If the process takes place in a fitness landscape/seascape *F*, the vector 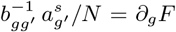 is its gradient, and *φ*(*x^t^*, *x*^*t*+Δ*t*^) is the rate at which the system climbs it up and therefore accumulates fitness (Fig. 1 and Eq. S8-S10 in [15]). This interpretation of fitness flux is exact in the diffusion approximation as Δ*t* → 0, but only approximate for the discrete formulas in Eq. (S105) and Eq. (S103).

We can take the expectation over *x*^*t*+Δ*t*^ to obtain the expected fitness flux per generation from any starting position *x^t^*,

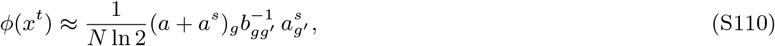

which is also plotted in Fig. S2A. Finally, we can take the expectation over *x^t^* to obtain the fitness flux bound in the diffusion approximation. It compares with the cost of selection bound as follows,

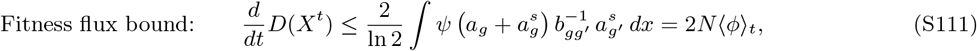

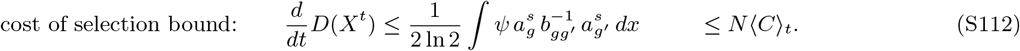

Note that in Eq. (S111) we added the factor 2 which was missing in [15], as pointed out in [23].

Again, the fitness flux bound requires the neutral process to be at a stationary distribution with detailed balance, which means zero neutral flux *u_g_* = 0 (see Eq. (S57)). (There is a more general flux theorem, which does not require detailed balance [15], but it does not provide a bound on Δ*D*(*X^t^*).) The single locus, two allele system satisfies the detailed balance, since diffusion only takes place along a single dimension. However, detailed balance is rare in systems with multiple loci with recombination and general forms of mutation.

#### S10C Comparisons of the discrete and the diffusion formulas

The two bounds, computed using both the Markov chain and the diffusion formulas, are compared in Fig. S2BC for the single locus, two allele system. Note that the bounds are obtained by averaging the functions 2*N_e_φ*(*x_A_*) and *N_e_C*(*x_A_*), such as those plotted in Fig. S2A, with respect to the distribution *ψ^X^* (*x_A_*).

In Fig. S2B, we use the Wright-Fisher model to compute the distribution over allele frequencies and the information increments. The population size is *N*_WF_ = *N_e_* = 100, mutation strength *Nμ* = 0.01 and selection strength varies across columns. The cost of selection bound is in black, and the diffusion formula (full line, based on Eq. (S112)) is in an agreement with the discrete formula (dashed line, based on Eq. (S104)). The discrete version of the fitness flux bound is violated, as the Wright-Fisher model does not satisfy detailed balance (grey dashed, based on Eq. (S103)). This is because cycles such as 0 → 1 → 2 → 0 copies of the *A* allele take place more often than the reverse cycle, since two alleles can get lost by drift in a single generation with a high probability, but are unlikely to arise by mutation. However, detailed balance holds in the diffusion approximation. If fitness flux is computed according to the diffusion formula, the bound holds under weak selection when the diffusion approximation is close to the Wright-Fisher model, but fails when selection is stronger (full grey line, based on Eq. (S111)).

In Fig. S2C, we use the Moran model of the same system to compute the distributions and information increment over time. Note that in order to have the same magnitude of genetic drift as the Wright-Fisher model, the Moran model needs twice as large a census population size, *N*_Moran_ = 2*N_e_*. Time is measured in generations (*N*_Moran_ = 2*N_e_* replacements each), and the increments in information are also computed per generation.

The discrete and diffusion formulas are again in agreement for the cost of selection bound. Note that we computed it as *N_e_*,〈*C*〉_*t*_ rather than *N*_Moran_〈*C*〉_*t*_, to account for the additional stochasticity due to random deaths, even though *N*_Moran_ = 2*N_e_* parents are sampled with selection in each generation. The fitness flux bound now holds in its discrete version – the Moran model only allows allele frequency changes by ±1/*N*_Moran_ and therefore satisfies detailed balance. If fitness flux is computed using the diffusion formula in Eq. (S111), it upper bounds the Moran model accumulation of information when selection is weak, but again fails when selection is strong and diffusion departs from the discrete model.

In conclusion, care is needed when applying the fitness flux bound to discrete models. In Main Text Fig. 4, we plot the information accumulation and the cost of selection bound based on the Wright-Fisher model, and the fitness flux bound based on the Moran model.

#### S10D Interpretation of the bounds under diffusion

The cost of control is non-negative and determined solely by the magnitude of the selection term 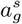, with the inverse drift covariance 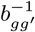, acting as the metric. As shown in Sec. S7 and Fig. S2A, this is proportional to the variance in fitness,

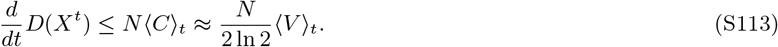

In Fig. S2BC, this bound starts off large and reduces slightly as selection removes variation from the population. It remains positive at the stationary state, where it represents the cost of maintenance.

In contrast, the fitness flux bound in Eq. (S111) contains the sum 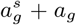, corresponding to selection and mutation contributions to the fitness flux. As a result, the fitness flux bound can be written as a sum of a selection term and a mutation term,

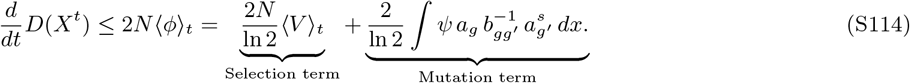

The selection term is proportional to the fitness variance – like the cost of selection bound, but with a 4 times higher numerical coefficient. When the selection term in Eq. (S114) dominates (a regime also discussed in [15]), the two bounds are proportional to each other, but the cost of selection bound is tighter. In general, the mutation term in Eq. (S114) causes the two bounds to behave in qualitatively different ways. Notably, the mutation term can be substantial and comparable to the selection term even when the parameters *N, s, μ* suggest that selection is strong and mutation is weak (i.e. under any or all of the conditions *Ns* ≫ 1, *Nμ* ≪ 1 and *s* ≫ *μ*). We illustrate and explain this in the following paragraphs.

Fig. S2A shows that the selection term in Eq. (S114) dominates especially at intermediate allele frequencies (around *x_A_* = 0.5), where the fitness variance is high and mutation in opposing directions cancels out. Near *x_A_* = 0, fitness variance is low and mutation towards the fitter allele *A* leads to a positive expected fitness flux. Near *x_A_* = 1, mutation towards the deleterious allele a dominates and leads to a negative expected fitness flux (Fig. S2A).

This enables a more detailed understanding of the fitness flux bound in the scenario in Main Text Fig. 4 and Fig. S2C. The system is initialized at the neutral stationary distribution, which is symmetric. Therefore the mutation term in fitness flux vanishes, because the positive and negative contributions (at *x_A_* < 0.5 and *x_A_* > 0.5 respectively) cancel out. The bound is therefore proportional to the average fitness variance. Over time, as selection shifts the distribution towards higher xA, mutation contributes more and more negatively, until the mutation and selection terms exactly cancel at stationarity. This happens regardless of *N, μ* and *s*. At stationarity, populations fluctuate around frequencies where mutation and selection are balanced, typically close to *x_A_* = 0 or *x_A_* = 1.

The only regime when the selection term dominates fitness flux for an extended period of time is when not only mutation is weak overall, but also the population is initialized at an intermediate frequency. This is shown in Fig. S3, which uses the same parameters as Fig. S2C, but the population is initialized at the frequency *x_A_* = 0.5.

Note that only the process with selection can be initialized at *x_A_* = 0.5. The neutral process must always be at the neutral stationary distribution to satisfy the assumptions of the fitness flux theorem. As a result, the information *D*(*X^t^*) is very high initially and decreases until drift spreads out the distribution *ψ^Xt^* towards *x_A_* =0 and 1. Later, *D*(*X^t^*) slightly increases over a much longer (mutation-limited) time scale, see Fig. S3A. The early phase is useful for illustrating the behavior of the fitness flux and the cost of selection, but neither bound is very informative there, as information is being lost.

In Fig. S3B we plot the increments in information, the fitness flux bound and the cost of selection bound per generation. Both bounds are proportional to the relative fitness variance in the early phase at intermediate frequencies, albeit with different proportionality constants (full and dotted purple lines in Fig. S3B). In the later phase, when the population is mostly fixed for one of the alleles, the mutation term in Eq. (S114) becomes important and negative, and the fitness flux bound departs from the fitness variance approximation.

### S11 Frequency dependent selection that maximizes fixation probability

In Main Text Fig. 3C,D, we compare the efficiency of selection (accumulated information per unit cost of selection) under constant selection, and under a specific form of frequency dependent selection. This frequency dependence is optimal in the sense that it maximizes the fixation probability of *A* at a given cumulative cost of selection. Below we describe the optimization procedure.

The calculation is done using a single locus, two allele Wright-Fisher model as described in Sec. S4, but instead of a single selection coefficient, we have a vector 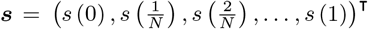 of selection coefficients *s*(*x_A_*) for each possible allele frequency *x_A_*. Given ***s***, we can compute

- The (right stochastic) transition matrix *P*(***s***) according to Eq. (S10), using the respective selection coefficient for each starting frequency (rows of *P*(***s***)).
- The vector *ψ*^fix^(***s***) of fixation probabilities for each possible starting frequency. It is given by the last column of the matrix power *P*(***s***)^*t*^ at infinite time *t* → ∞. Numerically, we keep doubling *t*, until the total probability that neither allele is fixed is less than 10^−6^ for all starting frequencies.
- The vector ***c***(***s***) of cost of selection at each frequency, 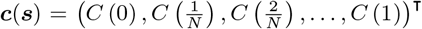. Note that *C*(0) = *C*(1) = 0 since there is no fitness variation when one of the alleles is fixed.
- The vector ***γ***(***s***) of expected total cost of selection until either allele is fixed, for each starting frequency. The first and last elements are again equal to zero, since one of the alleles is fixed and no more cost is incurred. The remaining elements can be computed using the recurrence relation

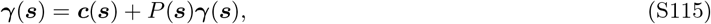

where ***c***(***s***) is the immediate cost and *P*(***s***)***γ***(***s***) is the expected future cost. Eq. (S115) can be solved for ***γ***(***s***) as a system of linear equations.
- The value vector ***ν***(***s***) = ***ψ***^fix^(***s***) – λγ(***s***), where λ is the Lagrange multiplier which quantifies the constraint on the total cost.

We look for ***s*** which maximizes value **ν**(***s***). This is an instance of a Markov decision process [24] similar to the pursuit/first passage problem [25] with a small modification to include an unwanted absorbing state (loss of the *A* allele). The frequency-dependent selection *s* corresponds to the decision policy. The optimal policy does not depend on time or the initial state, and maximizes all elements of ***ν***(***s***) simultaneously [25].

We optimize ***s*** iteratively. It is initialized at all zeros, and we alternate between a value update and a policy (selection) update,

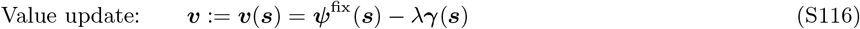

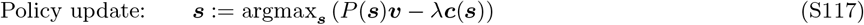

The value update uses the current estimate of ***s***. The policy update uses the current estimate of ***ν*** to compute the selection coefficient at each frequency, which maximizes the expected value in the next step, minus the immediate cost. The maximization is independent for each element of ***s*** and is done by binary searching for zero gradient (log_10_ s in range from 10^−5^ to 10^2^, binary search depth 10).

To produce Fig. 3C,D, we vary the cost constraint λ and compute ***s*** by 60 value-policy update iterations. Examples of the frequency dependent ***s*** are shown in Fig. S4A. Notably, selection is strongest at low frequencies of *x_A_* when *A* is at the greatest risk of being lost, but weak at higher frequencies to reduce costs. Fig. S4B,C show the fixation probability ***ψ***^fix^(***s***) and the expected total cost **γ**(***s***) for each starting frequency, from which the Main Text Fig. 3C,D uses only the values for the initial frequency 1/*N*.

**Figure S1:**
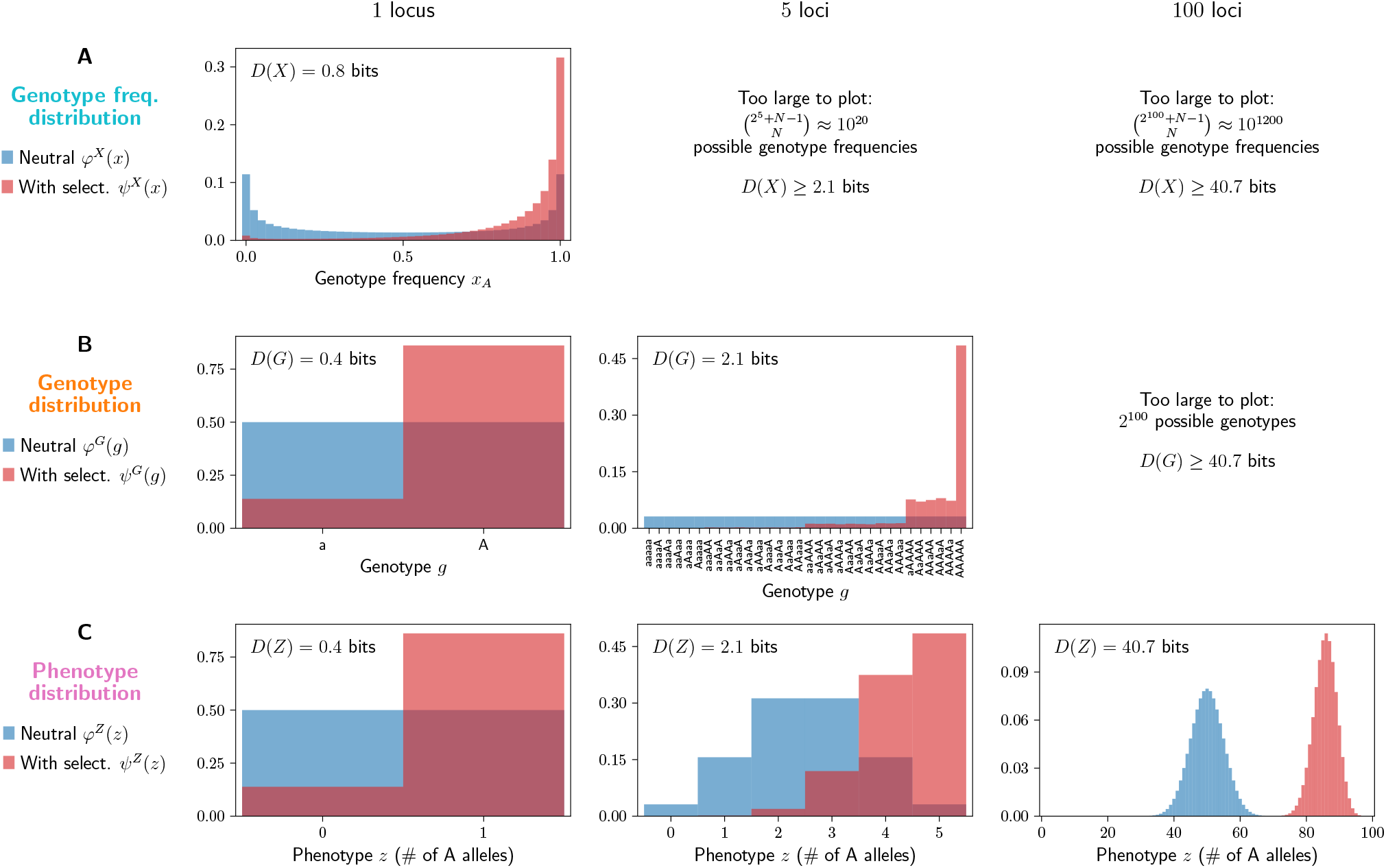
An example of the distributions over genotype frequencies, genotypes and phenotypes, with and without selection and for a varying number of loci, along with the corresponding measures of information. Based on a Wright-Fisher model with parameters *N* = 40, *μ* = 0.005, *s* = 0.05 (as in Main Text Fig. 1). Fitness is multiplicative across loci, *w* = (1 + *s*)^*z*^ where *z* is the number of beneficial *A* alleles carried – i.e. there is directional selection for an additive, fully heritable phenotype *z* (as in Main Text Fig. 6). All distributions are at the stationary state, computed using a transition matrix-based model (1 locus) or a long run (10^5^ generations after 200 generations of burn-in) of an individual-based model (> 1 locus). (A) Distributions over genotype frequencies (*ψ^X^* (*x*) with selection and *φ^X^* (*x*) without, red and blue) and the population-level information, *D*(*X*). The distributions cannot be plotted and *D*(*X*) cannot be directly estimated for 5 and 100 loci, due to the large number of possible genotype frequencies *x*, but we can still lower bound *D*(*X*) by *D*(*G*) or *D*(*Z*). (B) Distributions over genotypes (*ψ^G^*(*g*) with selection and *φ^G^*(*g*) without, red and blue) and the genotype-level information, *D*(*G*). This information is less than *D*(*X*), because selection not only gives preference to the fitter alleles, but also reduces the genetic variation within populations. The number of possible genotypes becomes too large for 100 loci, but we can still lower bound *D*(*G*) by *D*(*Z*). (C) Distributions over the phenotype (*ψ^Z^*(*z*) with selection and *φ^Z^*(*z*) without, red and blue) and the phenotype-level information, *D*(*Z*). The phenotype is simply the number of *A* alleles across all loci in an individual – an additive trait of varying polygenicity. In this example, selection favors the *A* allele at each locus, making fitness a function of *z*. In such cases, *D*(*G*) ≈ *D*(*Z*), since grouping genotypes into bins of equal *z* reduces the state space but preserves selective differences. In general, the trait might be unrelated to fitness or form only a component of it, leading to *D*(*G*) > *D*(*Z*). Note that *D*(*Z*) is approximately proportional to the number of loci, with each locus encoding about 0.4 bits. This is because loci evolve approximately independently, as there is zero epistasis, free recombination and little Hill-Robertson interference (see also Main Text Sec. 4B-C).

**Figure S2:**
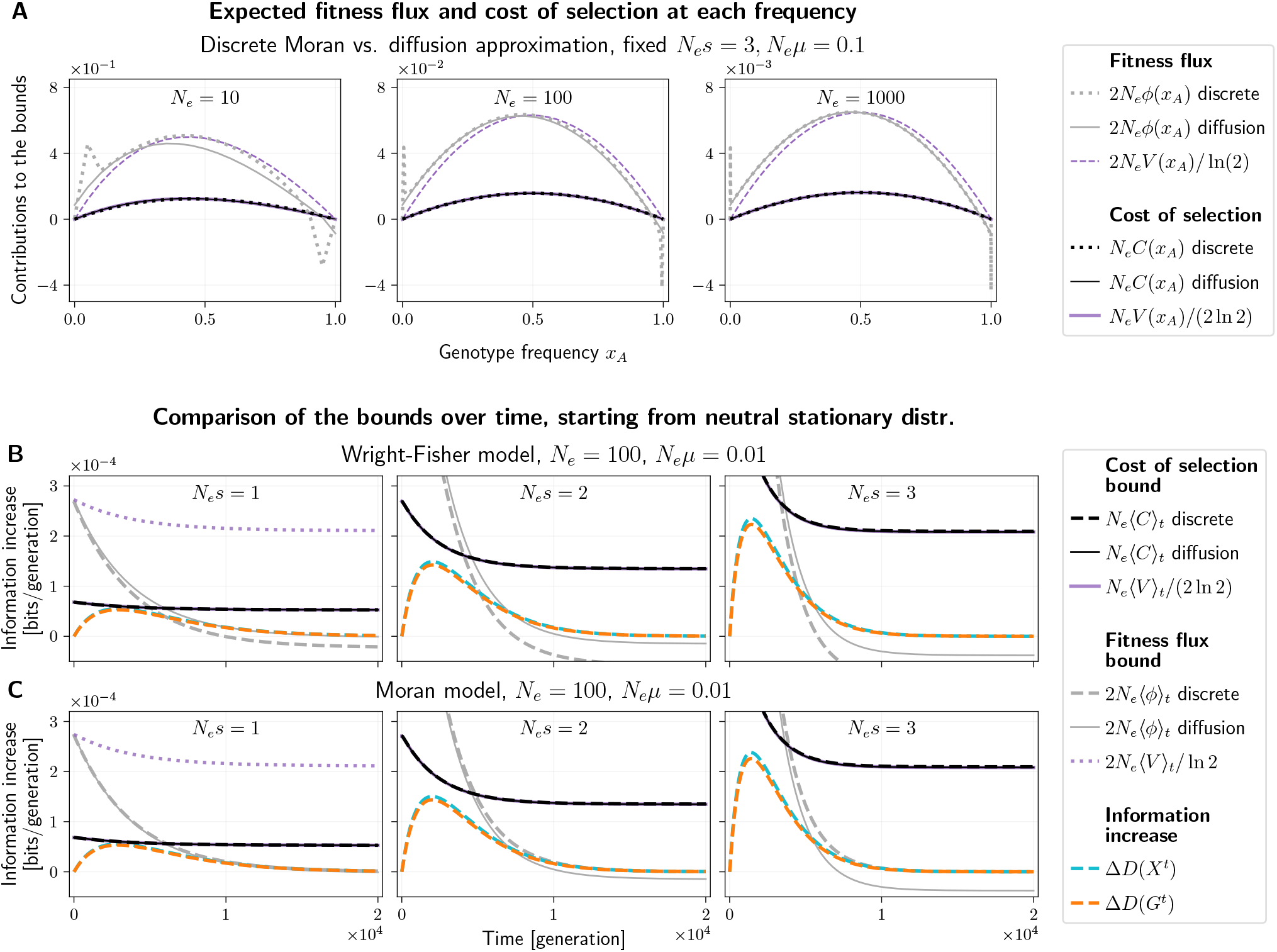
Comparisons between the fitness flux, the information-theoretic cost of selection and fitness variance, using the single locus, two allele systems under the Wright-Fisher model, the equivalent discrete-time Moran model and the diffusion approximation. (A) The expected fitness flux per generation (*ϕ*(*x_A_*), gray dotted) and the cost of selection (*C*(*x_A_*), black dotted) for each frequency *x_A_* under the discrete Moran-model for three different effective population sizes *N_e_* (left to right; census population size *N*_Moran_ = 2*N_e_* to account for the additional stochasticity in the Moran model compared to the Wright-Fisher model). We fixed *N_e_s* = 3 and *N_e_*μ = 0.1 to obtain models with a similar behavior but different time scales and granularity. Both *ϕ*(*x_A_*) and *C*(*x_A_*) are multiplied by *N_e_* and the same numerical factor as in the bounds on information accumulation rate (the bounds are obtained by averaging these values with respect to *ψ^X^* (*x_A_*)). The discrete formulas are compared with their diffusion approximation (full gray and black lines). As expected, the diffusion approximation is closer to the discrete model at higher *N_e_*. The fitness flux *ϕ*(*x_A_*) converges to the diffusion approximation non-uniformly across *x_A_*, due to the spikes next to the domain boundaries. These spikes do not disappear but get “squeezed out” at high *N_e_*. We also plot multiples of the relative fitness variance *V*(*x_A_*) (purple dashed and full lines), which approximate the fitness flux and the cost of selection. The cost *C*(*x_A_*) can be approximated very closely by *V*(*x_A_*)/(2 ln 2) as long as *s* ≪ 1 (here, the largest value is *s* = 0.3 for *N_e_* = 10). Fitness flux is the sum of a selection term proportional to *V*(*x_A_*) which is largest at intermediate *x_A_*, and a mutation term which dominates near *x_A_* = 0 or 1 and causes the discrepancy between *ϕ*(*x_A_*) and 2*V*(*x_A_*)/ ln 2. Even when mutation rate is small compared to selection, *μ* ≪ *s*, mutation is important as the system approaches the stationary distribution concentrated near *x_A_* = 0 and 1. (B) The Wright-Fisher model uses the same parameters as Main Text Fig. 4, namely *N* = *N_e_* = 100; *N_e_μ* = 0.01 and *N_e_s*, varied across columns. We plot the increase of information per generation (population level Δ*D*(*X*^*t*^), blue dashed; genotype level Δ*D*(*G^t^*), orange dashed), the upper bound in terms of cost of selection *N_e_*〈*C*〉^*t*^, computed using the discrete formula (Eq. (S104), black dashed) and the diffusion formula (Eq. (S112), black full), and the upper bound in terms of the fitness flux computed using the discrete formula (Eq. (S103), gray dashed) and the diffusion formula (Eq. (S111), gray full). We also show the fitness variance approximations of the two bounds (Eq. (S113), purple solid and the fitness variance term in Eq. (S114), purple dotted, outside the plot range for *N_e_s* = 2 and 3). The discrete fitness flux bound is violated, since the Wright-Fisher model does not satisfy detailed balance under neutrality. The continuous fitness flux bound holds under weak selection, but fails when selection gets stronger, as differences grow between the discrete system and the diffusion approximation. (C) The Moran model has the same effective parameters but double the census population size, *N*_Moran_ = 2*N_e_*. The curve descriptions are the same as for the Wright-Fisher model, but note that each generation consists of 2*N_e_* replacements, and the information increase as well as the upper bounds are rescaled accordingly. Also note that we plot the cost of selection bound *N_e_*〈*C*〉*t* using the effective population size *N_e_* rather than the census population size *N*_Moran_ = 2*N_e_*, which would lead to a twice as large (and less tight) bound. The Moran morel satisfies detailed balance under neutrality, and the discrete fitness flux bound holds under arbitrary selection. The continuous fitness flux bound again fails under strong selection. Note that the diffusion formula for the fitness flux bound would correctly upper bound the accumulation of information in a system modeled fully using the diffusion approximation. These figures show that it does not always upper bound the accumulation of information in discrete models, especially not when selection is strong. The Main Text Fig. 4 uses the information accumulation curves and cost of selection bounds based on the Wright-Fisher model, and the fitness flux bound based on the Moran model.

**Figure S3:**
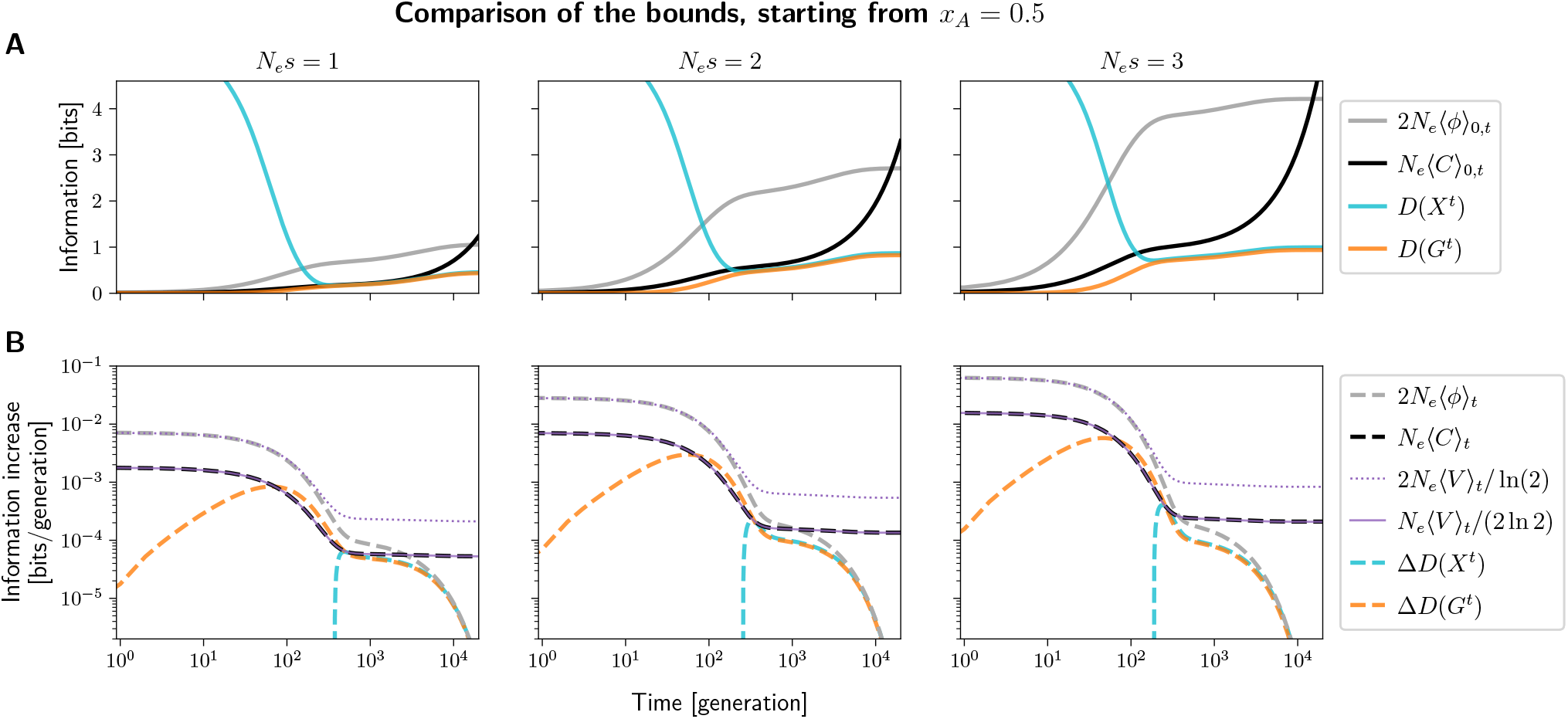
Demonstration of the fitness flux and the cost of selection bounds in the single locus, two allele system initialized at an intermediate frequency *x_A_* = 0.5. Note that only the process with selection is initialized at *x_A_* = 0.5; the neutral process must always be at the neutral stationary distribution to satisfy the assumptions behind the fitness flux theorem. This leads to a high initial value of *D*(*X^t^*) at *t* = 0. Calculated using the discrete Moran model with the same parameters as in Fig. S2C. (A) The cumulative information at the population level (*D*(*X^t^*), blue) and genotype level (*D*(*G^t^*), orange), as well the cumulative fitness flux (2*N_e_*〈*ϕ*〉_0, *t*_, gray) and cumulative cost of selection (*N_e_*〈*C*〉_0,*t*_, black). *D*(*X^t^*) starts out high due to the initial distribution being different under selection than under neutrality (*D*(*X*^0^) ≈ 12.3 bits, outside the plot range, in all three cases). At first, drift causes the distribution under selection to spread out towards extreme frequencies, *D*(*X^t^*) decreases on the time scale of ~ *N_e_* = 100 generations. Meanwhile, *D*(*G^t^*) accumulates as the mean frequency *x_A_* increases from 0.5 due to selection. The two measures of information eventually reach similar values, since mutation is low and the population is mostly fixed for one of the alleles. The cumulative fitness flux and cost of selection upper bound the cumulative information increase *D*(*X_t_*) – *D*(*X*^0^), but are not very informative in this case, because despite selection acting, *D*(*X^t^*) started at a value that is higher than can be maintained and is lost rather than accumulated. (B) The increase of information per generation on the population and genotype levels (Δ*D*(*X^t^*), blue dashed; Δ*D*(*G^t^*), orange dashed), and the upper bounds in terms of fitness flux (2*N_e_*〈*ϕ*〉_*t*_, gray dashed) and the cost of selection (*N_e_*〈*C*〉_*t*_, black dashed). Note the log scales on both axes. Initially, Δ*D*(*X_t_*) is negative and falls outside of the plot as drift spreads out the distribution over allele frequencies. Meanwhile, Δ*D*(*G^t^*) is positive as the mean frequency *x_A_* increases due to selection – this is associated with a positive fitness flux and cost of selection. After about 2*N_e_* = 200 generations, one of the alleles is likely to be fixed by drift, but the mean frequency continues to slowly increase at a mutation-limited rate (time scale 1/*μ* = 10^4^ generations), which also causes modestly positive Δ*D*(*X^t^*) and Δ*D*(*G^t^*). In this phase, the cost of selection remains nearly constant, while fitness flux slowly decays, providing a fairly tight bound on Δ*D*(*X^t^*). The cost of selection bound can be very well approximated with the relative fitness variance (*N_e_*〈*V*〉_*t*_/(2 ln 2), purple solid line). The fitness flux is proportional to the fitness variance in the first phase, when *x_A_* is near 0.5 (2*N_e_*〈*V*〉 / ln 2, purple dotted line), but departs from it later as *x_A_* tends to take values near 0 or 1 where mutation is important (see Fig. S2A). We note that similar behavior can also observed for different values of *N_e_μ, N_e_s*.

**Figure S4:**
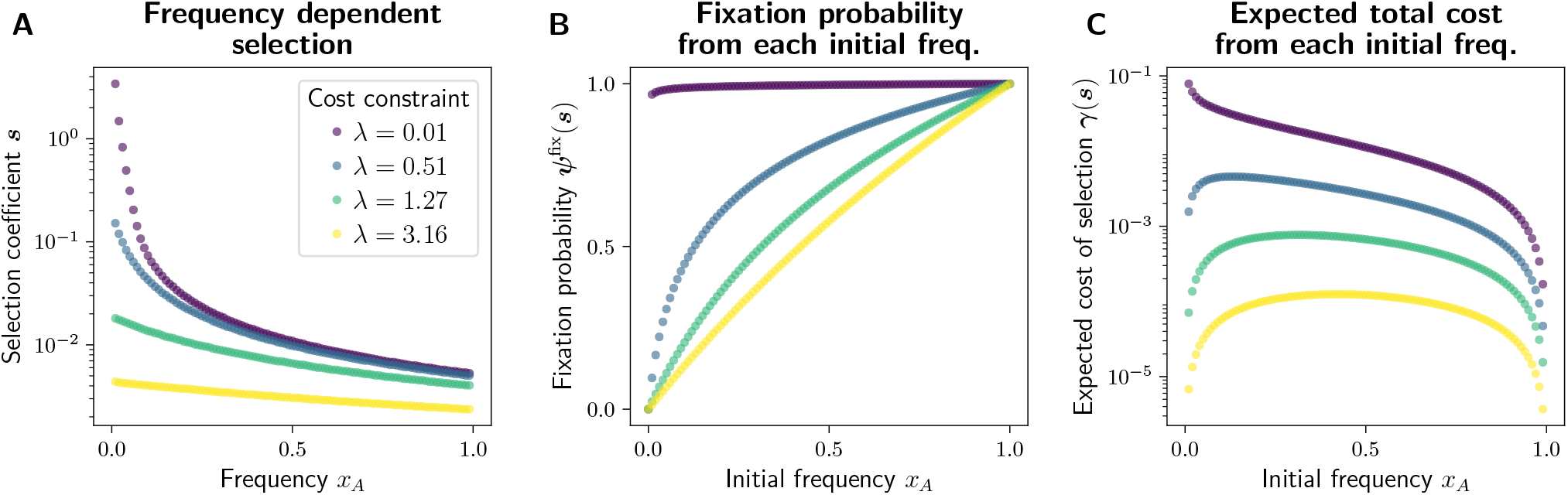
Frequency dependent selection that optimizes fixation probability of an allele at a constrained total cost. (A) The frequency dependent selection coefficient ***s***, computed as described in Sec. S11, under various cost constraints λ. When this constraint is greater, selection is weaker overall. (B) Fixation probability *ψ*^fix^(***s***) of the allele *A* for each starting frequency *x_A_*. The fixation probability is close to the frequency itself under weak selection (large λ), and higher when selection is overall stronger. (C) The expected total cost of selection, **γ**(***s***) associated with trajectories starting from each initial frequency *x_A_*. It is low for high frequencies, where the allele A is expected to be fixed soon, and for low frequencies under weak selection, where it is expected to be lost soon.

